# Comparative membrane proteomics reveals diverse cell regulators concentrated at the nuclear envelope

**DOI:** 10.1101/2023.02.13.528342

**Authors:** Li-Chun Cheng, Xi Zhang, Sabyasachi Baboo, Julie A Nguyen, Salvador Martinez-Bartolomé, Esther Loose, Jolene Diedrich, John R Yates, Larry Gerace

## Abstract

The nuclear envelope (NE) is a subdomain of the ER with prominent roles in nuclear organization, largely mediated by its distinctive protein composition. We developed methods to reveal novel, low abundance transmembrane (TM) proteins concentrated at the NE relative to the peripheral ER. Using label-free proteomics that compared isolated NEs to cytoplasmic membranes, we first identified proteins with apparent NE enrichment. In subsequent authentication, ectopically expressed candidates were analyzed by immunofluorescence microscopy to quantify their targeting to the NE in cultured cells. Ten proteins from a validation set were found to associate preferentially with the NE, including oxidoreductases, enzymes for lipid biosynthesis and regulators of cell growth and survival. We determined that one of the validated candidates, the palmitoyltransferase Zdhhc6, modifies the NE oxidoreductase Tmx4 and thereby modulates its NE levels. This provides a functional rationale for the NE concentration of Zdhhc6. Overall, our methodology has revealed a group of previously unrecognized proteins concentrated at the NE and additional candidates. Future analysis of these can potentially unveil new mechanistic pathways associated with the NE.

## Introduction

The nuclear envelope (NE) is a specialized sub-domain of the ER that forms the boundary of the nucleus and compartmentalizes chromosomes and associated metabolism (Dultz & Ellenberg, 2007). It is a double membrane sheet comprising the outer nuclear membrane (ONM), the inner nuclear membrane (INM) and the nuclear pore membrane connecting these (Obara *et al*, 2022). The pore membrane is juxtaposed to nuclear pore complexes (NPCs), transport channels spanning the NE that link the nucleus and cytoplasm (Beck & Hurt, 2017; Knockenhauer & Schwartz, 2016; Lin & Hoelz, 2019). NPCs are massive supramolecular assemblies (∼100 MDa in higher eukaryotes) containing multiple copies of ∼30 different proteins termed nucleoporins (or “Nups”). NPCs mediate nucleocytoplasmic trafficking of most macromolecules by facilitated mechanisms involving shuttling nuclear transport receptors that bind to their cargoes and specific Nups (Wing *et al*, 2022). NPCs also act as significant barriers to passive diffusion of molecules larger than ∼20-40 kDa (Beck & Hurt, 2017; Knockenhauer & Schwartz, 2016).

The ONM is contiguous with more peripheral ER and shares biochemical and functional properties with the latter, whereas the INM has nucleus-centered functions that are specified by its distinctive protein composition (Dultz & Ellenberg, 2007; Pawar & Kutay, 2021). In higher eukaryotes, the INM is lined by a polymeric meshwork of nuclear lamins, type V intermediate filament proteins (Burke & Stewart, 2013; Gruenbaum & Foisner, 2015). Three major lamin subtypes with discrete developmental expression profiles are found in vertebrates: lamins A/C, lamin B1 and lamin B2. Moreover, all eukaryotic cells contain a set of transmembrane (TM) proteins concentrated at the INM and at the nuclear pore membrane (Cheng *et al*, 2019; Pawar & Kutay, 2021; Schirmer *et al*, 2003). Collectively, nuclear lamins and these TM proteins have critical roles in the nuclear structure and mechanics (Cho *et al*, 2017; Maurer & Lammerding, 2019; Miroshnikova & Wickstrom, 2022), chromosome organization and maintenance (Hildebrand & Dekker, 2020; Kim *et al*, 2019), and regulation of signaling and gene expression (Choi & Worman, 2014; Gerace & Tapia, 2018). At least 15 human diseases arise from mutations in lamins and TM proteins of the NE, underscoring their functional significance (Shin & Worman, 2021; Wong & Stewart, 2020).

TM proteins of the INM become membrane-integrated in the peripheral ER during their synthesis. In higher eukaryotes, most of these proteins appear to accumulate at the NE by a diffusion-retention mechanism. This involves diffusive movement of the proteins in the lipid bilayer around NPCs, coupled with their binding to lamins, chromatin and/or other INM-associated components (Katta *et al*, 2014; Ungricht & Kutay, 2015). With this mechanism, exchange of TM proteins between ONM and INM is bidirectional and is limited by the size of their cytoplasmic domains (Katta *et al*., 2014; Ungricht & Kutay, 2015). By contrast, in yeast TM proteins commonly are transported to the INM by receptor-dependent facilitated diffusion around the NPC (King *et al*, 2006; Meinema *et al*, 2011), a mechanism that also contributes to INM targeting of some proteins higher eukaryotes (Mudumbi *et al*, 2020). The extent to which specific TM proteins accumulate at the NE relative to the peripheral ER is not fixed, but rather, can depend on the cell type (Malik *et al*, 2010) and its specific functional state (Le *et al*, 2016).

Most TM proteins concentrated at the NE are known to have specific functions at the INM and/or the NPC (Dultz & Ellenberg, 2007; Pawar & Kutay, 2021). Predicated on this logic, identification of novel NE-enriched proteins provides a framework to deepen an understanding of NE functions. To define such proteins, a comparative or “subtractive” proteomics approach has been deployed (Korfali *et al*, 2010; Schirmer *et al*., 2003; Tang *et al*, 2020; Wilkie *et al*, 2011), wherein isolated NEs are analyzed in tandem with purified microsomal membranes derived mainly from the peripheral ER, where most TM protein synthesis occurs. Proteins that are detected at higher levels in NEs than in microsomes by proteomics are candidate NE-concentrated proteins. However, this must be independently confirmed by immunolocalization and/or other methods.

These and allied approaches have identified a cohort of TM proteins concentrated at the NE, most of which are relatively abundant, i.e., present at more than ∼50-100,000 copies per nucleus (Cheng *et al*., 2019; Pawar & Kutay, 2021; Schirmer *et al*., 2003). However, the identification of low abundance NE-enriched proteins by these methods has been confounded by several technical issues. First, proteomics datasets reveal that the NE fractions contain well-defined TM markers of cytoplasmic organelles other than the ER, including plasma membrane, Golgi and mitochondria, indicating their partial co-fractionation with NEs (Korfali *et al*., 2010; Schirmer *et al*., 2003; Tang *et al*., 2020; Wilkie *et al*., 2011). Since non-ER cytoplasmic membranes are highly under-represented in the isolated microsomes used for comparative filtering (Korfali *et al*., 2010; Schirmer *et al*., 2003; Tang *et al*., 2020; Wilkie *et al*., 2011), some uncharacterized proteins that preferentially appear in the NE fraction may actually derive from other cytoplasmic organelles. Second, validation of NE-targeting of candidates has relied on ectopic overexpression (Korfali *et al*., 2010; Malik *et al*., 2010; Schirmer *et al*., 2003; Tang *et al*., 2020; Wilkie *et al*., 2011), often by transient transfection with chemical reagents. This can result in accumulation of ectopic proteins in non-physiological cellular aggregates that are often juxtanuclear (Tapia & Gerace, 2016), a pattern that can be confused with NE localization. Finally, in most cases quantitative evaluation of ectopic protein localization at the NE vs other cytoplasmic membranes was not rigorously implemented.

To circumvent these limitations, we developed modified proteomics-based methods for identification of low abundance proteins concentrated at the NE. Using chemically extracted membranes to enrich for TM proteins and thereby increase proteomics depth, candidates were identified by comparing isolated NEs to composite cytoplasmic membrane fractions rather than ER-biased microsomes. Subsequently, candidates enriched in the NE fraction were tested for selective targeting to the NE by low-level ectopic expression in cultured cells and systematic quantification by immunofluorescence microscopy. We evaluated this method by analyzing a cohort of NE-enriched TM proteins that we uncovered in a newly implemented comparative proteomics filtering (above), focusing on cell regulators not previously linked to the NE. We found that the majority of these candidates showed clear enrichment at the NE by immunofluorescence microscopy when expressed ectopically, confirming the NE association suggested by comparative proteomics.

We initiated a functional analysis of one of the low-abundance NE-enriched proteins identified in our screen, the palmitoyltransferase Zdhhc6 (Lakkaraju *et al*, 2012). We determined that a major NE palmitoylation target of Zdhhc6 is the INM-localized oxidoreductase Tmx4 (Cheng *et al*., 2019). Our analysis of palmitoylation-deficient mutants of Tmx4 suggested that Zdhhc6 modulates the NE concentration of Tmx4, providing a functional rationale for the observed NE-enrichment of Zdhhc6. Altogether, our methodology has revealed a cohort of previously unrecognized NE-enriched proteins and numerous additional candidates, many with interesting functional annotations not previously connected to the NE. This provides a robust framework for hypothesis development and testing to augment current functional understanding of the NE.

## Results

### Membrane Proteomics to Identify Candidate NE-concentrated Proteins

We sought to develop improved procedures to predict and validate TM proteins concentrated at the NE based on comparative proteomics with isolated NEs (Malik *et al*., 2010; Schirmer *et al*., 2003). Since most well-characterized NE proteins are relatively abundant (> 50-100,000 copies per nucleus), we wished in particular to uncover low abundance NE proteins. For this analysis we analyzed membrane fractions isolated from C3H10T1/2 murine mesenchymal stem cells (designated “C3H” cells below), a model for human diseases linked to the NE (Worman *et al*, 2010).

Since a major limitation of previous NE proteomics involved imprecise exclusion of proteins from cytoplasmic membranes that co-fractionated with the NE (see Introduction) (Korfali *et al*., 2010; Schirmer *et al*., 2003; Tang *et al*., 2020; Wilkie *et al*., 2011), we used membrane fractions comprising an ensemble of all cytoplasmic organelles, instead of ER-biased microsomal membranes, to filter out “background” (Fig. 2*A*).

Starting with a cell homogenate obtained by hypotonic cell lysis (sample 1), we prepared a low-speed nuclear pellet (sample 3), which also contained a substantial amount of trapped peripheral ER (∼50% of the total calnexin) and other cytoplasmic membranes. The postnuclear supernatant from this step (sample 2) was fractionated by flotation on a sucrose step gradient to yield light cytoplasmic membrane (LCM, sample 4) and heavy cytoplasmic membrane (HCM, sample 5) fractions. In parallel, the resuspended nuclear pellet (sample 3) was fractionated on a sucrose step gradient to yield a composite LCM/HCM fraction (designated HCM2, sample 6) and isolated nuclei (sample 7). The nuclear fraction then was used to isolate NEs (sample 9) by nuclease digestion and high salt extraction (Cheng *et al*., 2019).

To increase the proteomics depth for detection of TM proteins, we evaluated methods for enriching TM proteins in the isolated membranes. We compared the effects of two chemical perturbants known to preferentially deplete peripheral membrane proteins and cytoskeletal elements from membranes (Fig. 1*A*), 6 M urea (Foisner & Gerace, 1993; Steck & Yu, 1973) and 0.1 M Na_2_CO_3_ pH 11.5 (Fujiki *et al*, 1982). Initially we used Western blotting to monitor marker proteins in the NE and LCM fractions treated with these conditions (Fig. 1*B*). In NEs extracted with 6 M urea (“urea”), the peripheral membrane protein lamin A as well as the cytoskeletal proteins actin, vimentin and myosin were efficiently removed as compared to unextracted membranes, whereas the NE-enriched TM proteins LAP2*β* and emerin (Cheng *et al*., 2019) were not detectably depleted. By contrast, treatment of NEs with 0.1 M Na_2_CO_3_ (“carbonate”) did not discriminate between TM and non-TM proteins as efficiently, since emerin was partially extracted and lamin A was incompletely depleted. In LCMs, there was no detectable difference in the extraction resistance of marker TM proteins of mitochondria (Tim23) and ER (calnexin), or in the loss of actin (Fig. 2*B*) with both chemical conditions.

**Fig. 1.**
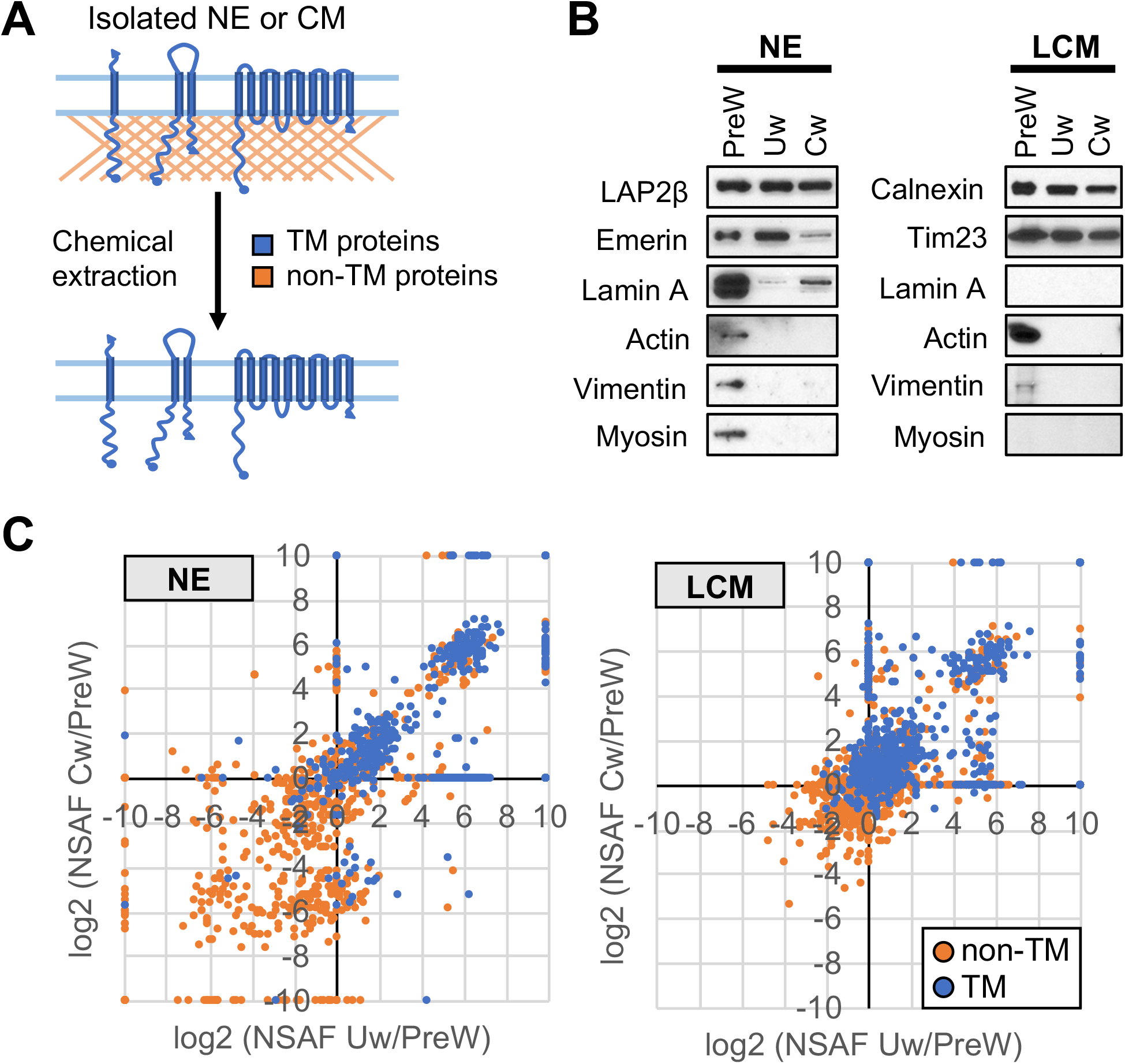
Evaluation of chemical extraction conditions for TM protein enrichment. *A,* Schematic diagram depicting the preferential solubilization of proteins lacking TM segments from membranes that is achieved with 6 M urea or 0.1 M Na_2_CO_3_. *B,* Western blot analysis of SDS gels to evaluate the presence of marker proteins (labeled on left of gels) in NE and LCM fractions. The experiment compared membranes without chemical extraction (PreW) to membranes following treatment with urea (Uw) or carbonate (Cw). *C,* Graphs comparing NSAF values of non-TM (brown symbols) and TM proteins (blue symbols) in NE and CML fractions in urea-treated (Uw) or carbonated-treated (Cw) membranes, as compared to their abundance without extraction (PreW). UniProtKB designations were used to define TM proteins. In the analyses in *B)* and *C)*, NE and LCM membrane fractions from a particular preparation were divided into three aliquots; one was used for PreW analysis, one for Uw and one for Cw.

**Fig. 2.**
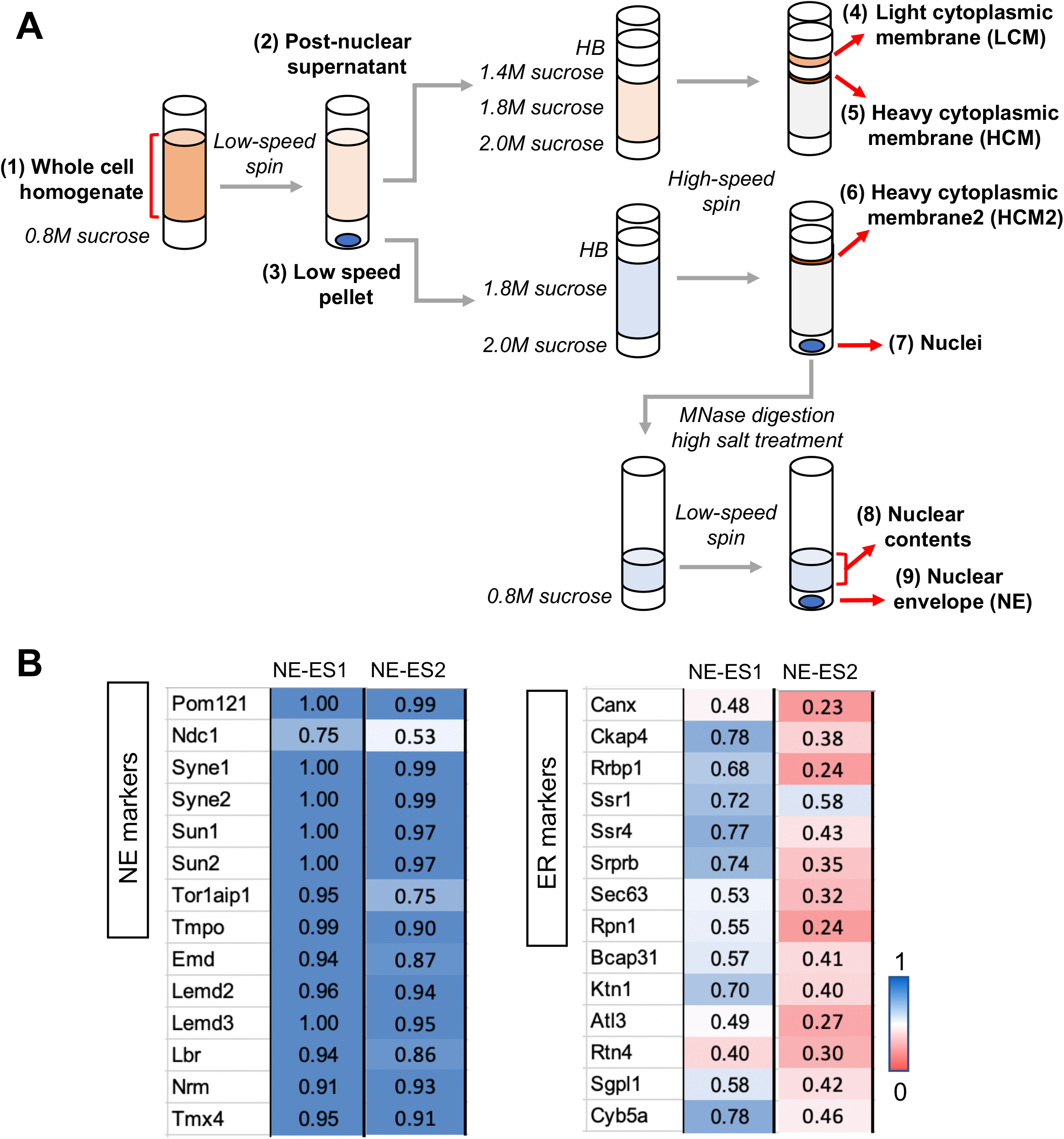
Isolation and evaluation of membrane fractions used for comparative proteomics. *A,* Schematic diagram depicting isolation of membrane fractions. The whole cell homogenate from C3H cells (1) was layered over a cushion containing 0.8 M sucrose and centrifuged at low speed to yield a post-nuclear supernatant (2) and a low speed pellet (3). Top: Sample (2) was adjusted to 1.8 M sucrose and introduced into a discontinuous sucrose gradient, above a layer containing 2.0 M sucrose, and below a layer with 1.4 M sucrose. The latter was overlaid with a zone of homogenization buffer (HB). Centrifugation at high speed resulted in appearance of a large opaque zone at the HB/ 1.4 M sucrose interface (LCM) and a smaller opaque zone at the 1.4 M sucrose / 1.8 M sucrose interface (HCM). Very little material was seen at the 1.8 M sucrose/ 2.0 M sucrose interface. The LCM and HCM fractions were harvested for subsequent analysis. Middle: Sample (3) was resuspended in buffer containing 1.8 M sucrose, and introduced into a discontinuous gradient above a layer containing 2.0 M sucrose and below a layer containing HB. Centrifugation at high speed yielded a pellet of isolated nuclei (7), and an opaque zone at the HB/ 1.8 M sucrose interface (6), which was harvested and designated HCM2. Bottom: Sample (7) was resuspended in HB, digested with micrococcal nuclease, adjusted to 0.5 M NaCl, and centrifuged at low speed over a cushion containing 0.8 M sucrose. The pellet from this step (9) comprised the NE fraction. *B,* Evaluation of NE-enrichment scores (NE-ES) in the NE fraction for marker TM proteins of the NE and ER. Values compare less stringent (NE-ES1) and more stringent (NE-ES2) calculation methods, as described in the text. Left panel, NE markers; right panel, ER markers. Ckap4, Rrbp1, Ssr1/4 and Ktn1 are enriched in sheet ER, whereas Atl3 and Rtn4 are more concentrated in tubular ER (see text).

We further compared these two conditions by analyzing NE and LCM fractions by MudPIT (mullti-dimensional protein identification technology) (Washburn *et al*, 2001), involving label-free LC/MS/MS (Table S1). We used normalized spectral abundance factor (NSAF) values to semi-quantitatively represent relative protein abundance (Zybailov *et al*, 2005). For each protein, the NSAF ratios in chemically extracted vs unextracted membranes for the NE (Fig. 1*C*, left panel) and LCM (Fig. 1*C*, right panel) fractions described the enrichment obtained with each chemical condition, and enabled comparison of the two methods (Tables S1 and S2). In a graphical depiction of these results (Fig. 1*C*), proteins whose relative abundance increased with both chemical extraction conditions appeared in the upper right quadrant, and comprised mainly TM proteins. Conversely, proteins with diminished abundance in both conditions were in the lower left quadrant and comprised mostly non-TM proteins. Both extraction conditions enriched TM proteins with strong preference over non-TM proteins. In the NE fraction (Fig. 1*C*, left), somewhat more TM proteins showed enrichment by urea but not carbonate extraction (lower right quadrant), as compared to enrichment by carbonate but not urea extraction (upper left quadrant). Considering these results together with the Western blotting (Fig. 1*B*), extraction of membranes with 6 M urea seemed to be more optimal than 0.1 M Na_2_CO_3_ for efficient recovery of TM proteins, and accordingly was deployed for subsequent analyses.

Altogether we carried out proteomics of NE and LCM fractions from four independent membrane preparations, analyzing samples extracted with urea from all four preparations (Table S1). In addition, we examined HCM and HCM2 fractions that were extracted with urea for three of the four preparations. To estimate protein abundance in NEs relative to other membrane fractions, we deployed composite NSAF data to compute NE enrichment scores (NE-ES) (Table S3). One metric, termed NE-enrichment score 1 (NE-ES1), incorporated all comparable proteomics data that was available for the four preparations. This involved NE and LCM fractions that were extracted with urea or carbonate or were not extracted (Table S3). For each protein, NE-ES1 is the ratio of the summated NSAF values from the NE fractions of these samples, compared to summated NSAF values from the (NE + LCM) fractions. We also calculated a second, more stringent, NE-enrichment score 2 (NE-ES2) for the three preparations extracted with urea, for which HCM and HCM2 data also was obtained. NE-ES2 is the ratio of summated NSAF values from the NE fractions, compared to the combined NSAF values for all fractions analyzed (NE, LCM, HCM and HCM2).

We evaluated the NE-ES1 and NE-ES2 scoring methods by considering abundant and well-characterized TM proteins of the NE and ER (Fig. 2*B,* see annotations in Table S3). For the NE markers, all proteins with the exception of Ndc1 had NE-ES1 values > 0.9. Moreover, the NE-ES2 values for these proteins were only marginally lower than NE-ES1, reflecting the minimal detection of these components in the HCM fractions. The lower NE-ES values for Ndc1 (0.75, NE-ES1; 0.53, NE-ES2) is most likely due to a substantial peripheral ER pool of this protein in C3H cells, as seen for emerin with certain cell types and conditions (Berk *et al*, 2013; Le *et al*., 2016).

A substantial number of the ER marker proteins had relatively high NE-ES1 values (∼0.7-0.8), particularly proteins reported to be enriched in sheet ER (Ckap4, Rrbp1, Ssr1/4, Ktn1) (Obara *et al*., 2022; Shibata *et al*, 2010). However, the NE-ES2 values for these proteins were markedly lower, in most cases < 0.5, due to the inclusion of HCM in the scoring calculation. These comparisons suggest that uncharacterized proteins with both NE-ES1 and NE-ES2 values > 0.8 are strong candidates for consideration as NE-concentrated species. The reliability of these metrics increases with the level of spectral detection for individual proteins, with more confident assessments involving detection across many or all of the fractionation experiments (Table S1). NE-ES1 incorporates more spectral data in this regard, although it is less stringent than NE-ES2.

It is important to note that NE enrichment scores are meaningful only for proteins containing a TM domain(s). This is because many intranuclear proteins that are undetectable in isolated cytoplasmic membranes, including components of chromatin and ribonucleoprotein particles, co-fractionate with NEs to a minor extent and are not quantitatively extracted with urea or carbonate treatment. Because they are absent from cytoplasmic membranes, these contaminants have high NE enrichment scores even though they are not concentrated at the NE *in situ*.

### Validation of candidate NE-concentrated proteins by quantitative immunofluorescence microscopy

We selected sixteen TM proteins identified from the proteomics screen, most with NE-ES1 and/or NE-ES2 scores > 0.9, to further evaluate as potential NE-concentrated proteins. For this we used a cell-based targeting assay involving immunofluorescence localization (Table 1, Fig. 3, Fig. S2). Fourteen of these (all except Tmem209 and Tmem214) were relatively low abundance based on their NSAF values, which were at least 5-10-fold lower than values for the TM nucleoporin markers Pom121 and Ndc1 (Fig. S1 and Table S1). Two members of this group, Tmem209 (Fujitomo *et al*, 2012; Malik *et al*., 2010) and Tmem53 (Korfali *et al*, 2011), were previously suggested to be NE-concentrated, but the remainder have not been characterized. The proteins had a range of sizes and predicted TM segments (Fig. S2). Categorized by annotated functions, they included enzymes for lipid biosynthesis and modification, oxidoreductases, signaling regulators and components involved in protein folding and related activities (Table 1).

**Fig. 3.**
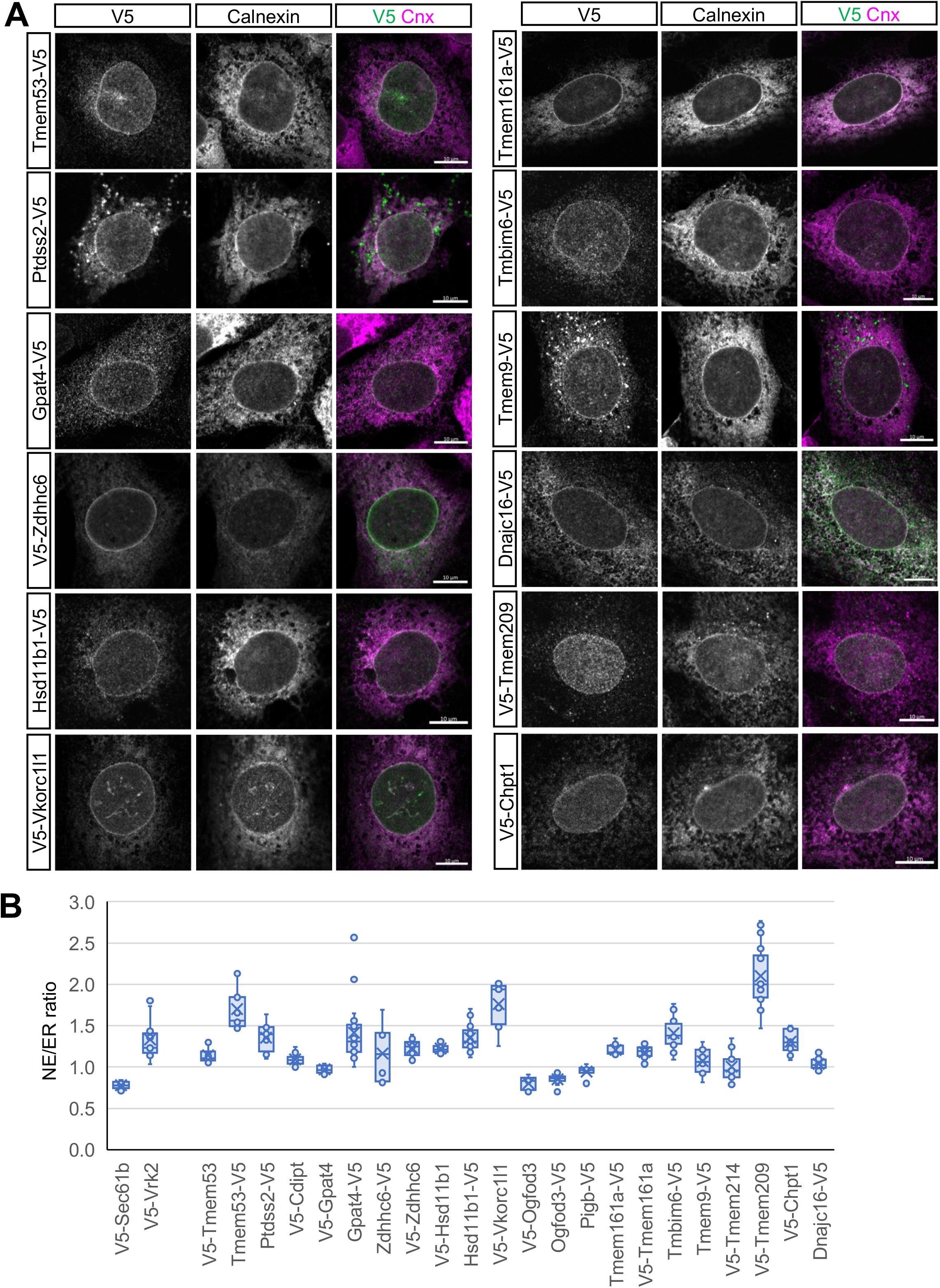
Immunofluorescence localization of candidate NE-enriched proteins by ectopic targeting assay. Proteins listed in Table 1 were tagged with a V5 epitope and transiently expressed in C3H cells using a lentiviral vector (see Fig. S2). Top panel: Montage of immunofluorescence images of representative cells expressing the indicated constructs, co-stained with antibodies that recognize the V5 epitope and endogenous calnexin. A merged image is shown in the third panel of each set. Bars, 10 μm. Bottom panel: Box and whisker plot depicting NE/ER concentration ratio for each construct, as determined by quantification of immunofluorescence images of at least 10 cells for each construct using the method depicted in Fig. S3. The NE/ER concentration of the control ER protein, V5-Sec61b, and of the previously identified NE-concentrated protein Vrk2 were determined in parallel.

**Table 1.**
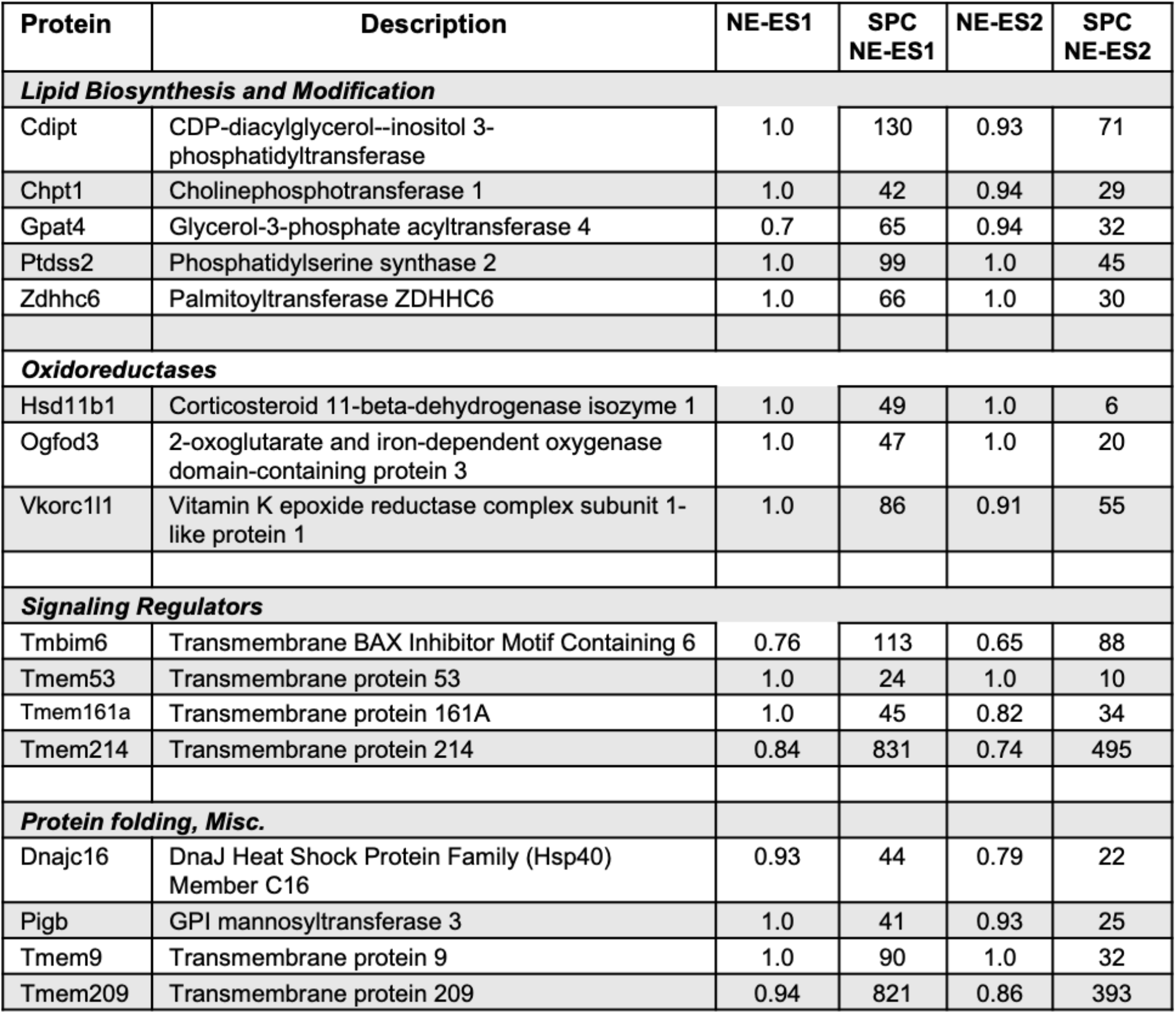
Proteins analyzed by immunofluorescence targeting assay.

We used a lentiviral vector and transient transduction conditions (Tapia & Gerace, 2016) to achieve low-level expression of the epitope-tagged candidates in C3H cells, and deployed immunofluorescence microscopy to compare the subcellular localization of ectopic V5-tagged constructs to endogenous calnexin, an abundant TM marker that is uniformly distributed throughout the NE/ER system (Shibata *et al*., 2010) (Fig. 3, see below). Low-level expression can be important for obtaining reliable NE targeting results, since some proteins that are endogenously concentrated at the NE are uniformly distributed throughout the NE/ER network or appear in large cytoplasmic inclusions with higher ectopic expression (Cheng *et al*., 2019). These effects, in part, likely reflect saturation of a limited number of binding sites for the proteins at the NE (Katta *et al*., 2014; Pawar & Kutay, 2021) and “spillover” of excess protein to contiguous ER elements.

For all 16 candidates analyzed, the ectopic constructs were localized to both the NE and peripheral ER, in a pattern broadly similar to calnexin (Fig. 3, top panel, Supplemental Fig. S4). Although none of the constructs appeared in large aggregates with our conditions, Tmem9 and Ptdss2 did occur in smaller, heterogeneous cytoplasmic foci as well as in ER-like structures. Visual inspection of the immunofluorescence patterns (Fig. 3, top panel) suggested that the majority of the candidates had a higher concentration at the NE than in the peripheral ER relative to calnexin. To analyze this quantitatively across populations of cells, we used a high information content method to calculate the relative concentration of ectopic constructs at the NE compared to the peripheral ER, designated the “NE/ER ratio” (Fig. S3). This method measured the fluorescence intensity of V5-tagged constructs (normalized to endogenous calnexin) in two circumferential bands around the nuclear periphery, one containing the NE and a second, more external band containing peripheral ER elements.

From this quantitative analysis, the majority of constructs (10 of the 16 proteins) had an average NE/ER concentration ratios of at least 1.2 (Fig. 3, bottom panel), indicating that these proteins indeed were concentrated at the NE with our expression conditions. For some proteins, a substantially higher NE/ER ratio was observed with the epitope tag on the N-vs C-terminus (notably Tmem53 and Gpat4). This may be explained by diminished epitope accessibility for the NE pool of the ectopic protein, or by tag-related inhibition of accumulation at the NE. Conversely, epitope tag placement had no effect on the NE/ER ratio for other proteins examined (Hsd11b1, Zdhhc6, Tmem161a). As a positive control, our quantification method yielded a NE/ER ratio of ∼1.3 for Vrk2, a previously described NE-concentrated protein (Birendra *et al*, 2017; Cheng *et al*., 2019). By contrast, the peripheral ER marker Sec61*β* appeared to be less concentrated at the NE than the adjacent ER (NE/ER, ∼0.8) as seen previously (Cheng *et al*., 2019), as were some of the candidates tested (Ogfod3, Pigb). We compared the NE targeting of Zdhhc6 and Hsd11b1 seen in transiently transduced C3H cells (Fig. 3) to that found in stably transduced C3H cells and in mouse embryonic fibroblasts (MEFs) (Fig. S4). We obtained similar targeting results with the different conditions, which all yielded NE/ER ratios of ∼1.2-1.4.

The NE/ER ratio calculated by the above method generally underestimates the actual *in situ* NE concentration of the ectopic proteins. This is because the circumferential band used to define the NE reflects the average intensity of all structures in this region. Limited by the resolution of light microscopy, this zone likely contains peripheral ER elements juxtaposed to the ONM and/or parts of the tubular elements connecting these two structures (Obara *et al*., 2022) as well as the NE itself. Moreover, since all well-characterized NE proteins are concentrated at specific sub-structural regions of the NE (Cheng *et al*., 2019; Pawar & Kutay, 2021; Schirmer *et al*., 2003), the average NE concentration of specific proteins reported by this method would underrepresent their concentration at discrete NE subdomains (e.g., INM or NPC).

Determination of the localization of the proteins described here with regard to NE substructure will require further analysis. However, the finely punctate labeling pattern on the nuclear surface seen for V5-Tmem209 was conspicuously reminiscent of NPC labeling (Fig. 3, upper panel, Fig. S5). Indeed, we observed strong co-localization of ectopic Tmem209 with the NPC-specific RL1 monoclonal antibody (Snow *et al*, 1987) (Fig. S5). Since Tmem209 is conserved in higher eukaryotes (PANTHER database) and its abundance is similar to that of the TM nucleoporins Pom121 and Ndc1 (Table S1), it may be a previously unrecognized TM nucleoporin.

In summary, the results from immunofluorescence localization of ectopically expressed constructs indicate that the majority of the candidates tested preferentially target to the NE relative to the peripheral ER with our conditions (see Discussion). Together with the proteomics data of membrane fractions, the results indicate that these proteins represent previously unrecognized NE-concentrated proteins.

### Functional analysis of the palmitoyltransferase Zdhhc6

An underlying prediction of this study is that some functions of NE-concentrated proteins are likely to involve the nucleus. To address this question, we carried out an initial analysis of the palmitoyltransferase Zdhhc6, one of the low abundance NE proteins identified here. Out of the seven palmitoyltransferases detected with more than 30 spectral counts in our analysis, Zdhhc6 was the only one with a high NE enrichment score. We determined that a pool of Zdhhc6 is localized to the INM in the vicinity of lamin B1 using the proximity ligation assay (Supplemental Fig. S6), indicating that it potentially could function at the INM. Even though we did not detect Zdhhc6 in LCM or HCM fractions by proteomics, a substantial peripheral ER pool of this protein seems likely (see Discussion). This is because Zdhhc6 has been implicated in palmitoylation of multiple peripheral ER resident proteins including calnexin (Lakkaraju *et al*., 2012), the IP3 receptor (Fredericks *et al*, 2014) and the ubiquitin E3 ligase gp78 (Fairbank *et al*, 2012) (see Discussion).

To identify potential NE palmitoylation targets of Zdhhc6, we depleted Zdhhc6 from C3H cells with shRNA and metabolically labeled the cells with the palmitic acid analog 17ODYA. (Fig. 4*A,* Fig. S7). After cell lysis, 17ODYA-containing proteins were tagged with biotin using click chemistry (Gao & Hannoush, 2018), modified proteins were captured on streptavidin beads, and samples were analyzed by tandem mass tag (TMT) proteomics to quantitatively compare protein bound to streptavidin beads with the different conditions (Table S4). Palmitoylated proteins were revealed by their selective or increased detection in samples incubated with 17ODYA as compared to palmitic acid. Moreover, palmitoylated proteins that were dependent on Zdhhc6 were reduced in the Zdhhc6 knockdown relative to control cells. This analysis detected five INM proteins whose capture levels were consistently reduced with Zdhhc6 knockdown (Fig. 4*B*, Table S4). We directly confirmed the palmitoylation of two of these proteins (lamin A and Tmx4), using Western blot analysis to detect biotinylated, 17ODYA-labeled proteins (Fig. 4*C*). Further, we found that the modification level of Tmx4 was reduced by over 50% in cells with knockdown of Zdhhc6 by siRNA (Fig. 4, *D* and *E,* Supplemental Fig. S7). No significant changes in 17ODYA labeling of lamin A were seen with our conditions, suggesting that lamin A palmitoylation was not limited by the reduced level of Zdhhc6 obtained with transient knockdown by siRNA, in contrast to the effects obtained with stable depletion by shRNA (Fig. S7 and Table S4). Together, our results indicate that Tmx4 is a palmitoylation target of Zdhhc6.

**Fig. 4.**
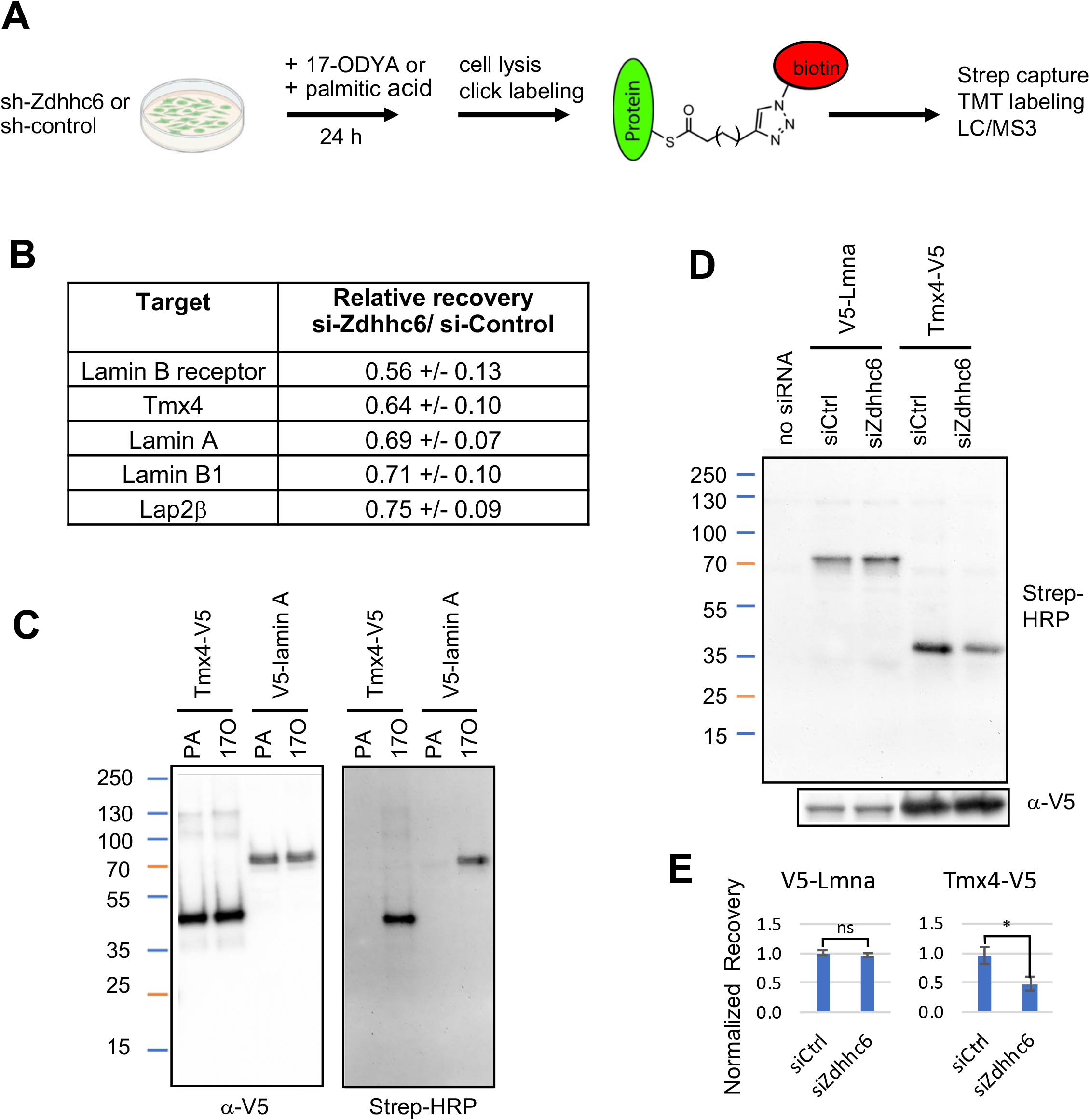
Identification of proteins palmitoylated by Zdhhc6 using metabolic labeling. *A,* Strategy used to identify proteins whose palmitoylation is dependent on Zdhhc6. C3H cells were stably transduced with an shRNA vector targeting Zdhhc6, or with either of two separate control shRNA vectors. Cells were incubated with either 17ODYA or palmitic acid for 24 hours, lysed and labeled by click chemistry to attach biotin to the alkyne group of 17ODYA. After complete solubilization with SDS, samples were incubated with streptavidin beads, which were washed and processed for on-bead digestion with trypsin. Eluted peptides were then labeled with TMT11 reagents and analyzed by LC/MS3. Two independent cell samples were prepared, and the TMT11 samples were analyzed in duplicate MS runs. *B,* List of NE proteins detected in all four repeats that showed a reduction in the level of bound protein with knockdown of Zdhhc6, indicating palmitoylation involving Zdhhc6. *C,* Confirmation of Tmx4 and lamin A palmitoylation. C3H cells that were stably transduced with vectors expressing Tmx4-V5 or V5-lamin A were incubated with 17ODYA (17A) or palmitic acid (PA) for 24 h, processed for click labeling with a biotin tag as in *(A)*, and analyzed by SDS PAGE followed by blot detection using streptavidin-HRP/*α*-HRP or *α*-V5 antibodies as indicated. *D,* Palmitoylation of Tmx4 and lamin A with Zdhhc6 depletion. C3H cells stably expressing Tmx4-V5 or V5-lamin A were treated with siRNA targeting Zdhhc6 or control siRNA. Cells then were incubated with 17ODYA or palmitic acid for 24 h, and processed with click labeling and analysis as in (*C)*. *E,* Quantification of changes in palmitoylation of Tmx4 and lamin A with knockdown of Zdhhc6 by siRNA, based on 2 repeats of the experiment shown in *(D).* Error bars represent SEM. * p < .05 (t test).

Protein palmitoylation has widely reported effects on protein localization and stability (Lakkaraju *et al*., 2012; Linder & Deschenes, 2007; Lynes *et al*, 2012). To evaluate whether palmitoylation similarly influences Tmx4, we first mapped Tmx4 palmitoylation sites. Calnexin, which is best characterized target of Zdhhc6, is modified on two cysteine residues positioned on the cytoplasmic side of the ER membrane, immediately adjacent to its TM segment (Dallavilla *et al*, 2016; Lakkaraju *et al*., 2012). Mouse Tmx4 contains two similarly positioned cysteine residues on residue 209 and 211, although only the Cys209 equivalent is conserved in human TMX4 (Fig. 5*A*). We made mutated versions of mouse Tmx4 in which either one or both of the conserved cysteine residues were mutated to alanine to block potential palmitoylation. Analysis of these constructs in cells labeled with 17ODYA revealed that most modification was lost with the C209A mutant, and that modification was virtually undetectable with the C209A/C211A mutant (Fig. 5*B*). This suggests that the conserved C209 on Tmx4 is the main site of palmitoylation and that C211 is modified to a lower extent.

**Fig. 5.**
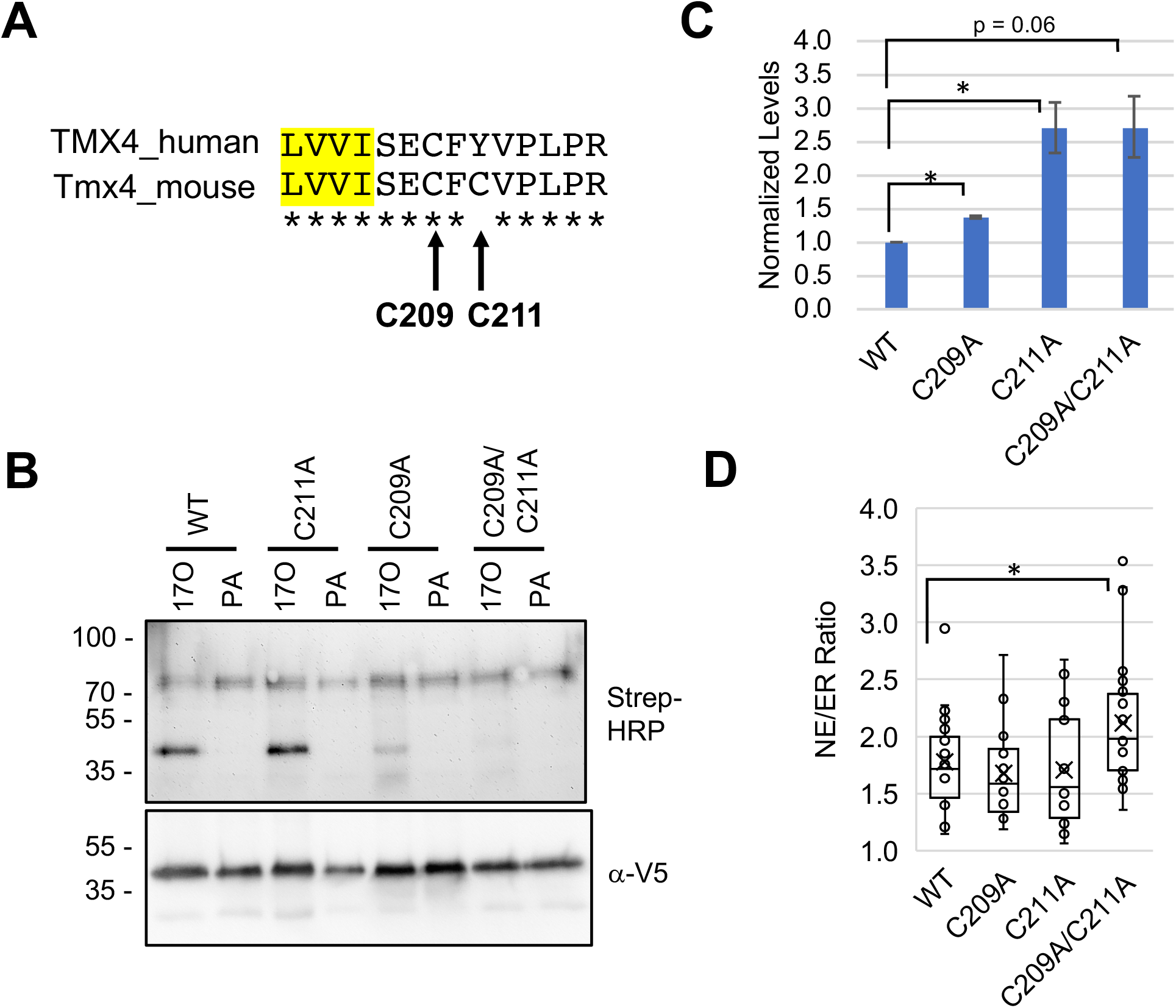
Sites of Tmx4 palmitoylation and palmitoylation effects on protein levels and localization. *A,* Sequence of human and mouse Tmx4 near potential palmitoylation sites. Yellow shading indicates the C-terminal end of the transmembrane domain, with the cytoplasmically oriented segment to the right. *B,* Palmitoylation in C3H cells stably transduced with V5-tagged versions of wild type (WT) Tmx4 or with the Tmx4 point mutants indicated. Cells were labeled for 24 h with either 17ODYA (17O) or palmitic acid (PA), processed by click labeling with a biotin tag and isolation on streptavidin-Sepharose, and finally analyzed by SDS PAGE with blot detection using streptavidin-HRP/*α*-HRP or *α*-V5 antibodies as indicated. Results similar to those shown were obtained in three experiments. *C,* Quantification of the level of V5-tagged Tmx4 constructs in stably transduced C3H populations with single-copy lentiviral integrations of the various constructs, using SDS PAGE and Western blotting with *α*-V5 antibodies. *D,* Quantification of the NE/ER concentration of various Tmx4 constructs in stably transduced C3H cell populations with single copy lentiviral integration. *C, D,* Error bars represent SEM. * p < .05 (t test).

Regarding the functions of Tmx4 palmitoylation, we found that levels of the C209A, C211A and C209A/C211A palmitoylation-deficient mutants were significantly increased in single-integration cell populations as compared to the WT protein (Fig. 5*C*). Moreover, the NE/ER concentration ratio was significantly increased for the C209A/C211A mutant as compared to wild-type Tmx4 (Fig. 5*D*). Together, these results suggest that palmitoylation both reduces the cellular levels of Tmx4 and diminishes its relative concentration at the NE. This points to palmitoylation as a means of controlling the levels of Tmx4 at the INM by two complementary mechanisms, and strongly suggests that Zdhhc6 has nucleus-related functions at the NE.

## Discussion

To promote an understanding of NE functions, we have refined methods to identify low abundance TM proteins that are concentrated at the NE. The core of our approach involved the use of label-free proteomics to analyze membrane fractions that were chemically extracted to enrich for TM proteins. By comparing isolated NEs to membrane fractions containing cytoplasmic organelles that partially co-fractionate with NEs, we identified proteins with selectively high NE abundance. To validate these candidates, proteins with high NE enrichment scores were ectopically expressed in cultured cells to quantify targeting to the NE from their site of synthesis in the peripheral ER by immunofluorescence microscopy.

The proteomics screen revealed numerous candidates with high NE enrichment scores (> 0.8) that were not previously connected to NE functions. Selecting a diverse sample set of mostly low abundance candidates for validation, we found that 10 of the 16 proteins examined targeted preferentially to the NE by immunofluorescence microscopy. Extrapolating from these results, it is likely that a substantial number of additional low abundance proteins with high NE enrichment scores also are enriched at the NE *in situ*. *A priori*, most of the NE-concentrated proteins we identified are likely to be concentrated at the INM and/or nuclear pore membrane, rather than the ONM (see Introduction).

Six of the candidates with high NE enrichment scores did not substantially concentrate at the NE in the ectopic targeting assay. It is possible that (some of) these were false positives that received high scores due to disproportionally low detection in the LCM and HCM fractions by MudPIT proteomics. Alternatively, they might indeed be NE-concentrated in C3H cells, but eluded detection due to several limitations intrinsic to the ectopic targeting assay. First, epitope accessibility might impair protein detection and/or targeting to the NE. This is exemplified by the considerably stronger NE concentration measured for the C-terminally tagged versions of Tmem53 and Gpat4 than the N-terminal versions. Second, ectopically overexpressed proteins might be undetectable above the levels that populate the peripheral ER if their NE accumulation is specified by a relatively low number and/or affinity of binding sites. Finally, the accumulation of certain proteins at the NE might require coordinated expression with other NE binding partners (e.g., other members of a multimeric complex) that is not reproduced with the ectopic conditions. Interestingly, two proteins with high NE enrichment scores that did not show NE concentration with our expression conditions, Tmem9 and Tmem214, were reported to interact with multiple NE proteins in stringent affinity capture-mass spectrometry screens with cultured cells (Hein *et al*, 2015; Huttlin *et al*, 2021). This is consistent with a NE association for these proteins that was not detected by our methods.

Whereas our proteomics and ectopic targeting methods identified proteins that are relatively concentrated at the NE vs the peripheral ER, a precise determination of their copy numbers in the two compartments *in situ* remains a formidable challenge. Genomic tagging of proteins (Cho *et al*, 2022) or use of specific antibodies in principle could address this issue, but both methods are currently limited by sensitivity for detection of low abundance proteins. Since the NE contains less than 5-10% of the surface area of the peripheral ER in most mammalian cells (Obara *et al*., 2022; Shibata *et al*., 2010), many NE-concentrated proteins may have a *greater copy number* in the peripheral ER than in the NE, despite their *selective concentration* at the NE. Viewed from this perspective, at least many of these proteins are likely to act in the peripheral ER, in addition to having NE-related functions (see below).

The protein set that we validated with the NE targeting assay spans multiple functional categories, including lipid biosynthesis, redox regulation and signaling control. Whereas the peripheral ER is a well-established site for lipid biosynthesis (Jacquemyn *et al*, 2017), the INM can be an important site for promoting synthesis of phosphatidylcholine and triglycerides under some conditions (Ohsaki *et al*, 2016; Romanauska & Kohler, 2018). Our findings that some phosphoglyceride synthesis isozymes are concentrated at the NE (Chpt1, Ptdss2, Gpat4) expands this current insight. Enrichment of Chpt1 at the INM is consistent with the finding that Pcy1a, the enzyme that synthesized CDP-choline and is rate-limiting for phosphatidylcholine synthesis, translocates from the nucleoplasm to the NE under conditions of phosphatidylcholine deficit (Cornell & Ridgway, 2015; Haider *et al*, 2018). Conceivably, the relative NE concentration of Chpt1 increases under similar conditions. Moreover, our observed NE concentration of Gpat4, the enzyme that catalyzes attachment of the first fatty acyl chain to glycerolphosphate (Jacquemyn *et al*., 2017), suggests that other downstream steps in phosphoglyceride biosynthesis may partly occur at the INM, in addition to the reactions catalyzed by Chpt1 and Ptdss2. Although the functional significance of lipid biosynthesis at the INM remains speculative, it can afford an mechanism to enhance coordination between the transcription of lipid biosynthetic genes and the end-products of their action (Jacquemyn *et al*., 2017).

Our detection of signaling regulators concentrated at the NE, including Tmbim6 and Tmem53, suggests a broader role for the NE in regulation of signaling than previously appreciated (Tapia & Gerace, 2016). Tmbim6 is a Bax-interactor that inhibits apoptosis in multiple cells models (Lebeaupin *et al*, 2020), and Tmem53 regulates cell cycle progression via several pathways (Korfali *et al*., 2011). Unraveling the mechanistic basis for these effects now can be considered in the context of the NE environment.

We previously found that the oxidoreductase Tmx4 is concentrated at the INM (Cheng *et al*., 2019), where it potentially could regulate the LINC complex and/or associated TorsinA, redox-sensitive proteins involved in connecting the nucleus to the cytoplasmic cytoskeleton (Cain *et al*, 2018; Lu *et al*, 2008; Zhu *et al*, 2010). Indeed, the oxidoreductase function of Tmx4 was recently implicated in disassembly of the LINC complex under pharmacological stress to induce NE autophagy (Kucinska, 2022). Our identification of Vkorc1l1 as an additional NE-concentrated oxidoreductase may expand this functional network since Vkorc1, a protein with a high NE enrichment score that is closely related to Vkorc1l1, interacts in a redox reaction with Tmx4 (Schulman *et al*, 2010). In a separate functional context, the NE enrichment of Hsd11b1, which reductively activates glucocorticoids and other steroids (Swarbrick *et al*, 2021), suggests a potential nuclear control system for positive regulation of steroid hormone action. Such a system also might include Steryl-sulfatase (Sts), another protein from our screen with high NE enrichment scores.

We focused on Zdhhc6 to investigate whether a low abundance protein concentrated at the NE could have nucleus-targeted functions. Our data indicated that several INM proteins including lamin A and the Tmx4 are its palmitoylation targets. Focusing on Tmx4, we identified the major sites of Tmx4 palmitoylation by Zdhhc6, and found that palmitoylation-deficient mutants were significantly increased in both steady state levels and in their NE/ER concentration ratio. Accordingly, palmitoylation of Tmx4 may attenuate Zdhhc6 functions at the INM both by decreasing its overall levels (most likely by increased proteosomal turnover (Dallavilla *et al*., 2016)) and by decreasing its INM concentration relative to the peripheral ER. This work strongly suggests a NE-selective function for Zdhhc6. Zdhhc6 itself is regulated by palmitoylation (Abrami *et al*, 2017), underscoring the potential regulatory complexity of this system.

In conclusion, the approaches we have outlined here, utilizing proteomic analysis of isolated membrane fractions and quantitative ectopic targeting assays, provide a template for confident identification of NE-concentrated proteins. Our findings have identified new NE-concentrated proteins that represent multiple functional categories, and have revealed a substantial number of additional candidates that remain to be directly evaluated. This information can provide a valuable framework for generating and testing new hypotheses on NE functions that heretofore have not been addressed.

## Materials and Methods

### Cell Culture

C3H/10T1/2 (C3H) cells (ATCC # CCL-225) and 293T cells (ATCC # CRL-3216) were acquired from the American Type Culture Collection and used at low passage number after freezing expanded stocks. Mouse embryonic fibroblasts (MEFs) were generated from C57BL/6 mice by immortalization with the SV40 T antigen. MEFs, C3H and 293T cells were maintained in high glucose Dulbecco’s modified Eagle’s medium (DMEM) (Gibco) supplemented with 10% fetal bovine serum (FBS) (Gibco), 1% penicillin/streptomycin/glutamine cocktail (Gibco), and 1% minimum essential medium non-essential amino acids (NEAA) (Gibco). All cells were maintained at 37°C in 5% CO_2_.

### Subcellular Fractionation and Membrane Extraction

For subcellular fractionation, C3H cells were seeded in 500 cm^2^ plates and allowed to reach 90% confluency. Plates were rinsed three times with ice-cold PBS, and then three times with ice-cold homogenization buffer (HB) (10 mM HEPES pH 7.8, 10 mM KCl, 1.5 mM MgCl_2_, 0.1 mM EGTA, 1 mM DTT, 1 mM PMSF, and 1 μg/ml each of pepstatin, leupeptin, and chymostatin). After these washes, cells were incubated in HB for 15 min on ice. Cells then were scraped off plates and were further disrupted by dounce homogenization with 18-20 strokes to achieve > 95% cell disruption. The whole cell homogenate was layered on top of 2 ml shelf of 0.8 M sucrose in HB and centrifuged at 2000 rpm for 10 min at 4°C in a JS5.2 rotor (Beckman Coulter) to yield a low speed nuclear pellet and a postnuclear supernatant, the latter comprising the zone above the sucrose shelf. The low speed nuclear pellet was resuspended in 1.8 M sucrose in HB using a cannulus, and the postnuclear supernatant was adjusted to a final concentration of 1.8 M sucrose in HB. The resuspended low speed nuclear pellet and the postnuclear supernatant were each layered in separate ultra-clear 13.2 ml nitrocellulose centrifuge tubes on top of a 1 ml layer of 2.0 M sucrose in HB. For the nucleus gradient, HB was layered over the loading zone to fill the nitrocellulose tube. For the postnuclear supernatant gradient, 1 ml of 1.4 M sucrose in HB was layered on top of the loading zone, followed by HB to fill the tube. The gradients then were centrifuged at 35,000 rpm (210,000g) for 1 h at 4°C with no brake in an SW41Ti rotor (Beckman Coulter). Two samples were collected from the nucleus gradient: a fraction comprising the HB/1.8 M sucrose interface, termed “heavy cytoplasmic membranes 2” (HCM2), and the pellet at the bottom of the 2.0 M sucrose layer, which comprised the purified nuclei fraction. The latter was collected by resuspension in HB and dounce homogenization with 2 strokes to disperse aggregates. For the postnuclear supernatant gradient, the HB/1.4 M sucrose interphase was collected and saved as “light cytoplasmic membranes” (LCM) and the 1.4 M sucrose/1.8 M sucrose interface was collected and saved as “heavy cytoplasmic membranes” (HCM). To prepare NEs, resuspended purified nuclei were incubated with 1 mM CaCl_2_ and 100 ku/ml micrococcal nuclease (New England Biolabs) in HB for 37°C for 15 min. Digested nuclei were then placed on ice and NaCl was added to a final concentration of 500 mM. The sample then was layered on top of 1 ml shelf of 0.8 M sucrose in HB and centrifuged at 4000 rpm for 10 min at 4°C in a JS5.2 rotor. A sample comprising the region above the 0.8 M sucrose layer was collected and saved as “nuclear contents”. The pellet was collected with resuspension in HB and saved as the NE fraction.

Chemical extraction of isolated membrane fractions involved four separate cell/membrane preparations. Isolated membrane fractions were dounce homogenized briefly and the protein concentration of each fraction was adjusted 0.5 mg/ml based on the Pierce BCA assay (ThermoFisher). To carry out the chemical extractions, 200 μl of each membrane fraction was added to 1.8 ml of the following solutions: deionized water (PreW, pre-wash), 100 mM sodium carbonate pH 11.5 (Cw, carbonate wash), or 8 M urea in 25 mM NaCl, 20 mM Tris-HCl pH 8.8 (Uw, urea wash). For the Cw, samples were incubated for 30 min at 4°C. For the Uw, samples were incubated 30 min at room temperature. Samples were centrifuged for 1 h at 75,000 rpm (239,000 g) at 4°C with a TLA100.3 fixed angle rotor. Pellets were gently rinsed with ice-cold deionized water and then processed for proteomics. Two membrane preparations were divided and used for parallel carbonate and urea extractions, and an additional two membrane preparations were used for urea extraction only.

### MudPIT Proteomics and NE Enrichment Scoring

MudPIT proteomics (Wolters *et al*, 2001) was carried out on NE and LCM fractions obtained from all four of the membrane preparations, and on HCM and HCM2 fractions obtained from three of these preparations. 30 µg protein from each subcellular fraction, estimated by the BCA protein assay, was suspended in 4 M urea, 0.2% RapiGest SF (Waters Corporation) and 100 mM NH_4_HCO_3_ pH 8.0. Proteins were reduced with Tris(2-carboxyethyl)phosphine hydrochloride (TCEP) and alkylated with 2-Chloroacetamide. Next, proteins were digested with 0.5 µg Lys-C (Wako) for 4 h at 37°C, and then for 12 h at 37°C in 2 M urea, 0.2% RapiGest SF, 100 mM NH_4_HCO_3_ pH 8.0, 1 mM CaCl_2_ with 1 µg trypsin (Promega). Digested proteins were acidified with TFA to pH < 2 and RapiGest SF was precipitated out. Each fraction was loaded on individual MudPIT micro-columns (2.5 cm SCX: 5 µm diameter, 125 Å pores; and 2.5 cm C18 Aqua: 5 µm diameter, 125 Å pores; Phenomenex), and resolved across an analytical column (15 cm C18 Aqua: 5 µm diameter, 125 Å pores) (Phenomenex).

Analysis was performed using an Agilent 1200 HPLC pump and a Thermo LTQ-Orbitrap Velos Pro using an in-house built electrospray stage. MudPIT experiments were performed with steps of 0%, 10%, 20%, 30%, 50%, 70%, 80%, 90%, 100% buffer C and 90/10 % buffer C/B (Wolters *et al*., 2001), being run for 5 min at the beginning of each gradient of buffer B. Electrospray was performed directly from the analytical column by applying the ESI voltage at a tee (150 mm ID) (Upchurch Scientific) (Wolters *et al*., 2001). Electrospray directly from the LC column was done at 2.5 kV with an inlet capillary temperature of 325°C. Data-dependent acquisition of tandem mass spectra were performed with the following settings: MS/MS on the 20 most intense ions per precursor scan; 1 microscan; reject unassigned charge state and charge state 1; dynamic exclusion repeat count, 1; repeat duration, 30 sec; exclusion list size 500; and exclusion duration, 90 sec.

Protein and peptide identification was done with the Integrated Proteomics Pipeline – IP2 (Integrated Proteomics Applications, Inc. http://www.integratedproteomics.com/). Tandem mass spectra were extracted (monoisotopic peaks) from raw files using RawConverter (He *et al*, 2015) and were searched against the UniProt SwissProt *Mus musculus* database (release 2018_01) with reversed sequences using ProLuCID (Peng *et al*, 2003; Xu *et al*, 2015). The search space included all fully-tryptic and half-tryptic peptide candidates with static modification of 57.02146 on cysteines. Peptide candidates were filtered using DTASelect (Cociorva *et al*, 2007) at 1% protein level False Discovery Rate, (parameters: -p 1 -y 1 --trypstat --pfp 0.01 --extra --pI -DM 10 --DB -- dm -in -t 0 --brief –quiet) (McDonald *et al*, 2004; Tabb *et al*, 2002).

The NSAF values for each protein in the various membrane fractions analyzed were used to calculate NE enrichment scores. NE enrichment score 1 (NE-ES1) utilized data from NE and LCM fractions obtained for all 4 membrane preparations and for both urea and carbonate extraction conditions. It involved the following equation, where e = experimental run and p = protein ID:

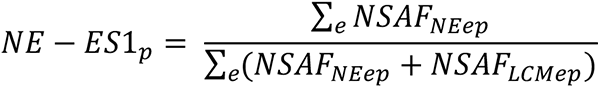

NE enrichment score 2 (NE-ES2) utilized the additional data for HCM and HCM2 fractions that was obtained from three of these preparations according to the following:

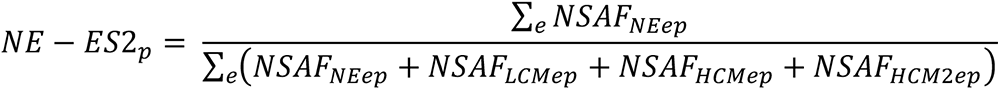

### Isolation of Palmitoylated Proteins by Metabolic Labeling

C3H cell populations were stably transduced with a lentiviral vector that produced either a) shRNA targeting Zdhhc6 (“shZdhhc6”), b) a control shRNA sequence not found in the mouse genome (“shCtrl”) or c) a second control shRNA targeting GFP (“shGFP”). Cells were used for metabolic labeling within 5 passages after lentiviral transduction/puromycin selection. Two biological replicates of the three cell populations were each analyzed with two LC/MS3 technical repeats. For each replicate, 5 x 10^5^ cells per 10-cm dish were seeded one day prior to metabolic labeling. On the day of labeling, cells were rinsed with PBS and 10 ml of growth medium containing 50 μM 17-Octadecynoic Acid (17ODYA) or 50 μM phosphatidic acid was added. After 24 h, cells were harvested by trypsinization and pelleting, and were washed twice with cold PBS. Cells then were resuspended in non-denaturing lysis buffer (20 mM Tris-HCl pH 8, 300 mM NaCl, 2 mM EDTA, 1% NP-40, supplemented with 10 μM Palmostatin B (Merck #17851) and Halt phosphatase and protease inhibitor cocktails (ThermoFisher #78440)) and were sonicated on ice for 2 cycles of 10 seconds with a Sonic Dismembrator 60 (ThermoFisher). The lysate was clarified by centrifugation at 4°C for 15 min at 20,000 g.

The clarified lysate then was precipitated with methanol-chloroform (4:1) to remove free 17ODYA, and 1 mg of protein from each sample was click-labeled with Biotin-PEG3-azide (Cayman) (0.58 mM Biotin-PEG3-azide, 1.15 mM TCEP, 1.15 mM CuSO_4_, 1X Tris((1-benzyl-4-triazolyl)methyl)amine solution in 1:4 DMSO/butanol) by incubating for 1 h at room temperature. Free biotin-PEG3-azide was removed by methanol-chloroform precipitation and the protein pellets were resuspended in 6 M urea supplemented with 2% SDS in PBS. Protein samples were reduced for 20 min with 5 mM TCEP and then were alkylated with 20 mM iodoacetamide in the dark. Samples then were diluted 10-fold with PBS and were incubated with 100 μl of a 50% streptavidin-Sepharose (Cytiva) bead slurry for 1.5 h at room temperature, washed four times with 2 M urea in PBS containing 0.2% SDS pH 7, four times with 2 M urea in PBS pH 7, and finally washed once with 50 mM tetraethylammonium bromide (TEAB). Protein-bound streptavidin-Sepharose beads were then stored at -80°C until further processing.

### TMT Sample Preparation, Data Acquisition and Analysis

Frozen streptavidin-Sepharose beads were resuspended in 100 mM TEAB, 8 M urea, reduced with 5 mM TCEP and alkylated with 10 mM 2-chloroacetamide. The bead suspension was diluted to 2 M urea, supplemented with 1 mM CaCl_2_ and digested with 2 μg trypsin (Promega) for 15 h at 37ᵒC. Beads were pelleted by centrifugation at 20,000 g for 5 min, and 150 μg of protein was removed from the supernatant. After pooling 1/10^th^ of each sample to create a reference, each sample was labeled with 11-plex TMT (ThermoFisher) at 3:1 (TMT:protein) ratio in 30% acetonitrile, according to the manufacturer’s recommendation. After labeling, the samples were pooled and acetonitrile was removed using vacuum centrifugation. The mixture of TMT-labeled peptides was acidified and fractionated using a high pH reversed-phase peptide fractionation kit (ThermoFisher) according to the manufacturer’s recommendation. The fractions were dried by vacuum centrifugation and resuspended in 5% acetonitrile, 0.1% formic acid.

TMT-labeled peptides were analyzed using an EASY-nLC 1200 UPLC coupled with an Orbitrap Fusion Lumos mass spectrometer (ThermoFisher). LC buffer A (0.1% formic acid, 5% acetonitrile in H2O) and buffer B (0.1% formic acid, 80% acetonitrile in H_2_O) was used for all analyses. Peptides were loaded on a C18 column packed with Waters BEH 1.7 μm beads (100 μm x 25 cm, tip diameter 5 μm), and separated across 180 min: 1-40% B over 140 min, 40-90% B over 30 min and 90% B for 10 min, using a flow rate of 400 nL/min. Eluted peptides were directly sprayed into MS via nESI at ionization voltage 2.8 kV and source temperature 275 °C. Peptide spectra were acquired using the data-dependent acquisition (DDA) synchronous precursor selection (SPS)-MS3 method. Briefly, MS scans were done in the Orbitrap (120k resolution, automatic gain control AGC target 4e5, max injection time 50 ms, m/z 400-1500), the most intense precursor ions at charge state 2-7 were then isolated by the quadrupole and CID MS/MS spectra were acquired in the ion trap in Turbo scan mode (isolation width 1.6 Th, CID collision energy 35%, activation Q 0.25, AGC target 1e4, maximum injection time 100 ms, dynamic exclusion duration 10 s), and finally 10 notches of MS/MS ions were simultaneously isolated by the orbitrap for SPS HCD MS3 fragmentation and measured in the Orbitrap (60k resolution, isolation width 2 Th, HCD collision energy 65%, m/z 120-500, maximum injection time 120 ms, AGC target 1e5, activation Q 0.25).

Protein and peptide identification was done with the Integrated Proteomics Pipeline – IP2 (Integrated Proteomics Applications, Inc. http://www.integratedproteomics.com/). Tandem mass spectra were extracted (monoisotopic peaks) from raw files using RawConverter (He *et al*., 2015) and were searched against a UniProt SwissProt *Mus musculus* database (#UP000000589), including streptavidin (UniProt #P22629), with reversed sequences and standard contaminants, using ProLuCID (Peng *et al*., 2003; Xu *et al*., 2015). The search space included all fully-tryptic and half-tryptic peptide candidates with static modification of 57.02146 Da on cysteines and 229.1629 Da on lysines and N-termini. Peptide candidates were filtered using DTASelect (Cociorva *et al*., 2007) at 1% spectrum level false discovery rate, (parameters: -p 2 -y 1 --trypstat --fp 0.01 --extra --pI -DM 10 --DB --dm -in -t 1 --brief –quiet) (McDonald *et al*., 2004; Tabb *et al*., 2002).

### Molecular Cloning

To construct lentiviral vectors expressing V5-tagged versions of our proteins of interest, the following cDNA clones were purchased from Origene: pCMV6-Chpt1 (RefSeq NM_144807), pCMV6-Pigx, (NM_024464)and pCMV6-Dnajc16 (NM_172338). The remaining genes were cloned from a cDNA library constructed from the C3H cells. Genes of interest were amplified by PCR using Q5 High-Fidelity DNA Polymerase (New England Biolabs #M0491L). All genes were inserted into either pLV-Ef1a-V5-LIC-IRES-PURO (Addgene #120247) or pLV-Ef1a-LIC-V5-IRES-PURO (Addgene #120248). pLV-EF1a-IRES-Puro LIC-compatible vectors were digested with SrfI (New England Biolabs). PCR fragments and SrfI-digested vector were combined with NEBuilder HiFi DNA Assembly Master Mix (New England Biolabs #E2621) in a 2:1 ratio of insert to vector. The DNA and DNA Assembly Master Mix were incubated at 50°C for 20 min and then NEB Stable Competent Cells (New England Biolabs) were transformed with the product. Cells were incubated at 30°C for 24 h, and resulting colonies were picked for clone validation. All cDNA clones were confirmed by complete DNA sequencing of the ORF in both 5’-3’ and 3’-5 directions. All clones for lentiviral vectors expressing the proteins described in this study are available from Addgene.

### Lentiviral transduction of cells

To produce lentiviruses, 293T cells at 60-80% confluency were shifted to DMEM supplemented with 10% FBS, 1% glutamine, and 1% NEAA without antibiotics 30 min prior to transfection. Cells were transfected with pRSV-REV (gift from Didier Trono, Addgene plasmid #12253), pMDL-RRE (gift from Didier Trono, Addgene plasmid #12251), pCMV-VSVg (gift from Bob Weinberg, Addgene plasmid #8454), and a pLV-EF1a-gene-of-interest vector (pLV-EF1a-GOI) using Lipofectamine 2000 (Invitrogen). Viral supernatant was harvested 48 h after transfection and filtered through a 0.45 μm polyethersulfone membrane filter (GE Healthcare Whatman). Western blotting of the viral supernatants with anti-V5 antibodies (below) was used to assess expression of the encoded V5-tagged constructs.

For stable lentiviral transduction of C3H cells and MEFs, cells were diluted to 5 x 10^4^ cells/ml in DMEM with 10% FBS, 1% glutamine, and 1% NEAA without antibiotics, and polybrene (EMD Millipore) was added to a final concentration of 10 mg/ml. Cells were transduced with different viral loads (ranging from 1 to 500 μl of viral supernatant per 1 ml of cells) to obtain cell populations with a range of multiplicities of infection. After 3 days, cells were treated with puromycin (Invitrogen) to select for cells that had integrated viral DNA. C3H cells and MEFs were treated with 5 μg/ml and 1 μg/ml puromycin, respectively, for up to 1 week. Cell populations with < 30% cell survival under puromycin treatment, reflecting mostly single-integration events, were further expanded and grown for Western blotting and immunofluorescence. Transient lentiviral transduction of C3H cells involved the methods described above, except cells were analyzed 48 h after treatment with virus without puromycin selection.

### Depletion of Zdhhc6 by RNAi

Analysis of the palmitoylation targets of Zdhhc6 by proteomics utilized C3H cell populations that were stably transduced with lentiviral vectors producing shRNA, using the methods described above. Cells with depletion of Zdhhc6 were transduced with sh-Zdhhc6 (Dharmacon #RMM3981-201743749). Control cells were transduced with sh-eGFP (Dharmacon #RHS4459) and non-targeting control shRNA (Dharmacon #RHS6848) (see Fig. 4). Levels of Zdhhc6 mRNA in the stable populations were determined by qRT-PCR as described (Cheng *et al*, 2022).

Subsequent experiments with Zdhhc6 depletion involved the use of C3H populations that had been stably transduced with various V5-tagged Tmx4 lentiviral constructs. Zdhhc6 depletion in these cells was achieved with ON-TARGETplus siRNA SmartPool oligonucleotides, one targeting Zdhhc6 (Horizon Discovery #L-059510-01-0005) and a second comprising a non-targeting control pool siRNA (Horizon Discovery #:D-001810-10-05). For this procedure, siRNAs were dissolved in 60 mM KCl, 6 mM HEPES pH 7.5, 0.2 mM MgCl_2_ and added to serum-free DMEM to a concentration of 50 nM. This solution was then diluted to a final concentration of 25 nM siRNA with a 1:4 mixture of Dharmafect-1 (DF-1) reagent (Horizon Discovery #T-2001-03) and DMEM, respectively, and was incubated at room temperature in the dark for 20 min. Using C3H populations grown to a density of 5 x10^5^ cells per 10-cm plate, the siRNA-DF-1 mixture was added drop-wise to cells to a final concentration of 5 nM and distributed evenly with gentle rocking. After growth for 24 h, the medium was replaced with DMEM supplemented with serum and antibiotics. At 48 h post-transfection, cells were analyzed by Western blotting and qRT-PCR (Cheng *et al*., 2022).

### Antibodies

The following primary antibodies/reagents were used for Western blotting: rabbit anti-calnexin (Sigma #C4731), rabbit anti-lamin A (affinity purified, made in-house to residues 391-428 of human lamin A), rabbit anti-LAP2*β* (made in-house to residues 1-194 of rat LAP2*β*), mouse anti-Tim23 (BD Transduction Laboratories #611222), streptavidin-HRP (Thermo Fisher S911), mouse anti-actin (clone C4, gift from Dr. Velia Fowler), mouse anti-myosin heavy chain (clone 3J14, US Biological # M9850-15B), mouse anti-vimentin (clone RV202, Abcam # ab8978-1). The secondary antibodies used for Western blot detection were: sheep anti-mouse HRP (GE Healthcare #NA931), donkey anti-rabbit HRP (GE Healthcare #NA934V).

The following primary antibodies were used for immunofluorescence staining: mouse anti-V5 (Invitrogen #46-0705), rabbit anti-V5 (ThermoFisher #PA1-993), rabbit anti-calnexin (Abcam #ab22595), guinea pig anti-lamin A/C (made in-house to gel-purified rat liver lamin A), mouse monoclonal IgM RL1 (recognizing FG repeat nucleoporins (Snow *et al*., 1987)), mouse anti-nesprin-1 (8C3, gift from Dr. Glenn E. Morris, RJAH Orthopaedic Hospital, UK), mouse anti-nesprin-1 (7A12, Millipore #MABT843), rabbit anti-nesprin-2 (ThermoFisher, Invitrogen #46-0705), rabbit anti-nesprin-3 (United States Biological Corporation), rabbit anti-myosin heavy chain (Abcam #124205). DAPI (Sigma) was used to stain DNA.

### Western blotting

For Western blotting, cells were resuspended in 2X Laemmli buffer (4% SDS, 10% 2-mercaptoethanol, 20% glycerol, 0.004% bromophenol blue, and 0.125 M Tris-HCl pH 6.8) and boiled for 5 min. Samples were run on a Novex Tris-Glycine gel (Life Technologies) using FASTRun Buffer (Fisher Scientific). Samples were then transferred to a nitrocellulose membrane (Life Technologies). Membranes were rinsed twice with Tris-buffered saline (TBS) with 0.1% Tween-20 (Tw) and then blocked with 5% bovine serum albumin (BSA) in TBS/Tw. Membranes were incubated with primary antibody diluted in 0.5% BSA in TBS/Tw overnight at 4°C. Membranes were then washed 6 times with TBS/Tw and incubated with HRP conjugated secondary antibodies in TBS/Tw for 1 h at room temperature. Signals were then developed using an enhanced chemiluminescence kit (Thermo Fisher) for 5 min before exposure to film.

### Immunofluorescence Staining

For immunofluorescence staining, cells were plated on sterile glass coverslips the day before analysis. 24 h after plating, cells were rinsed with PBS containing calcium and magnesium and fixed using 2% paraformaldehyde (PFA) (Electron Microscopy Sciences # 15710) in PBS for 20 min. Samples were rinsed three times with PBS and blocked for 15 min using PBS with 5% goat serum (Jackson ImmunoResearch Laboratories) and 0.5% Triton X-100 (Tx) (Fisher Scientific). Samples were then incubated with primary antibody diluted in PBS with 1% goat serum and 0.1% Tx overnight at 4°C. After washing with PBS/0.1% Tx four times, samples were incubated with Alexa Fluor conjugated secondary antibody diluted in PBS/0.1% Tx at room temperature for one h. Samples were finally washed twice with PBS/0.1% Tx, incubated with DAPI at room temperature for 10 min, and then washed twice with PBS and mounted on glass slides using Aqua-Poly Mount (Polysciences).

The proximity ligation assay was carried out as described previously (Cheng *et al*., 2022), using C3H cells that had been stably transduced with a lentiviral vector expressing Myc-lamin B1.

### Light Microscopy and Quantification

Confocal images were acquired on a Zeiss 780 or a Zeiss 880 Airyscan laser-scanning confocal microscope with a 63X PlanApo 1.4 NA objective. Contrast adjustment of the representative images was performed with ZEN software (Zeiss). 10 or more images from each stably or transiently transduced cell population of the lowest expression levels were randomly chosen and the NE/ER ratio was quantified. Lamin A staining was used to outline the nucleus and the area of NE and ER were defined by -0.5 to 0 μm (NE) and +0.5 to +1 μm (ER) relative to the lamin A-defined nuclear edge using the “*Enlarge*” function in ImageJ (NIH). Total fluorescent intensities of V5 staining in both areas were measured and normalized to the calnexin staining of the same area. The ratio of NE/ER was then calculated by dividing the normalized V5 signals in the NE to the normalized V5 in the ER.

The co-localization analysis was performed with the “*Coloc2*” function in ImageJ. Where necessary, raw images was processed using the rolling-ball “*background subtraction*” function in ImageJ. Control and test images were processed with identical parameters. Representative images were prepared with automatic Airyscan processing in ZEN.

## Data Availability

The proteomics datasets have been deposited in the public proteomics repository MassIVE (Mass Spectrometry Interactive Virtual Environment), part of the ProteomeXchange consortium (Vizcaino *et al*, 2014), with the identifier MSV000091154 and is available through the following link: https://massive.ucsd.edu/ProteoSAFe/private-dataset.jsp?task=417b3a6d8c4e4809b247dfcbb728de32.

## Acknowledgements

The authors thank Dr. Scott Henderson for advice on confocal microscopy, Titus Jung for assistance with proteomics data management and Gerace and Yates lab members for helpful discussions.

## Author contributions

L.G. conceptualization; L-C. C., S. B., and X. Z. data curation;

L-C. C. and S. M-B. formal analysis; L-C. C., X. Z., J. N., S. B., and J. D. investigation; L-C. C. methodology; L.G. project administration; E.L., J. N., and J. Y. resources; L-C. C. and L.G. supervision; L-C. C. and L.G. visualization; L.G. writing – original draft; L.G writing – review & editing; L.G. and J. Y. funding acquisition

## Abbreviations

The abbreviations used are:

NE: nuclear envelope
TM: transmembrane
ONM: outer nuclear membrane
INM: inner nuclear membrane
NPC: nuclear pore complex
MEF: mouse embryonic fibroblast
DMEM: Dulbecco’s modified Eagle’s medium
FBS: fetal bovine serum
NEAA: nonessential amino acids
HB: homogenization buffer
LCM: light cytoplasmic membranes
HCM: heavy cytoplasmic membranes
MudPIT: multidimensional protein identification technology
17ODYA: 17-Octadecynoic Acid
TMT: tandem mass tag
NSAF: normalized spectral abundance factor
NE-ES: nuclear envelope enrichment score

## Supplemental Figure Legends

**Fig. S1.**
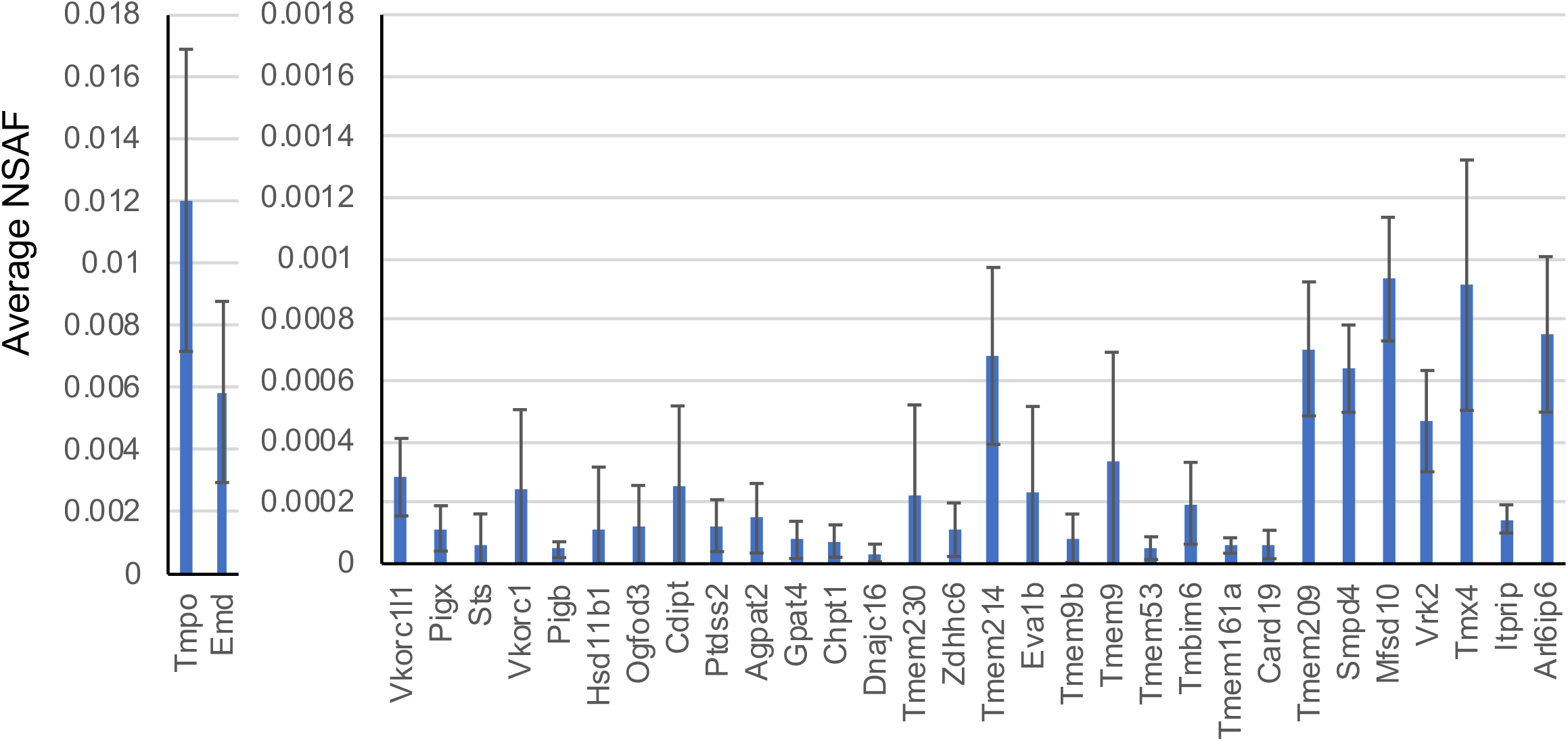
Estimation of the relative abundance of proteins analyzed in this study by NSAF values. Graph represents composite data from all four cell fractionation experiments. Error bars indicate SEM.

**Fig. S2.**
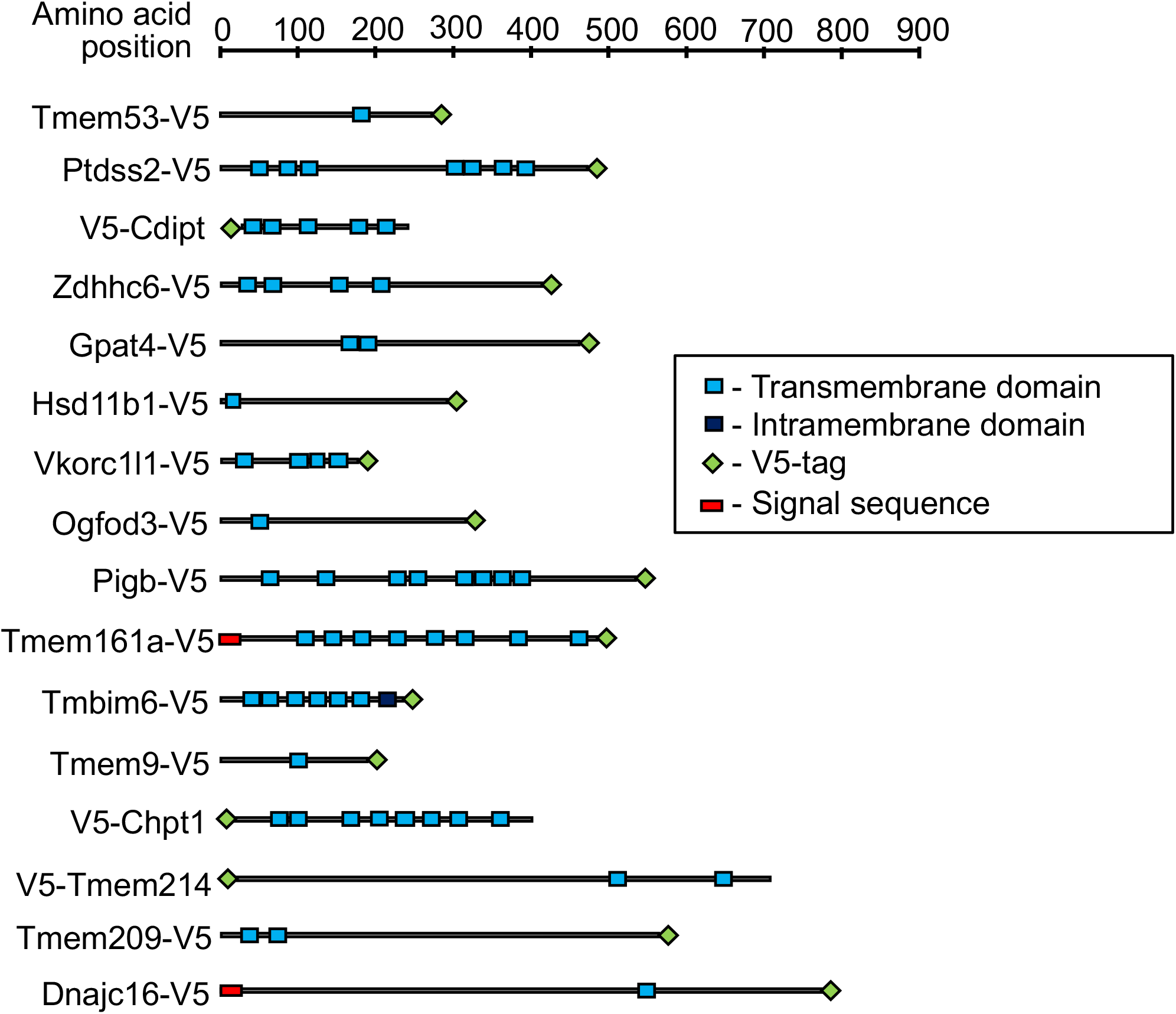
Schematic diagram depicting the sizes, transmembrane segments and epitope tag placement of proteins analyzed in this study. The work also analyzed versions of Tmem53, Gpat4, Zdhhc6, Hsd11b1, Ogfod3 and Tmem161a with an N-terminal V5 epitope tag (not shown), in addition to the diagrammed constructs with a C-terminal tag. Error bars represent SEM.

**Fig. S3.**
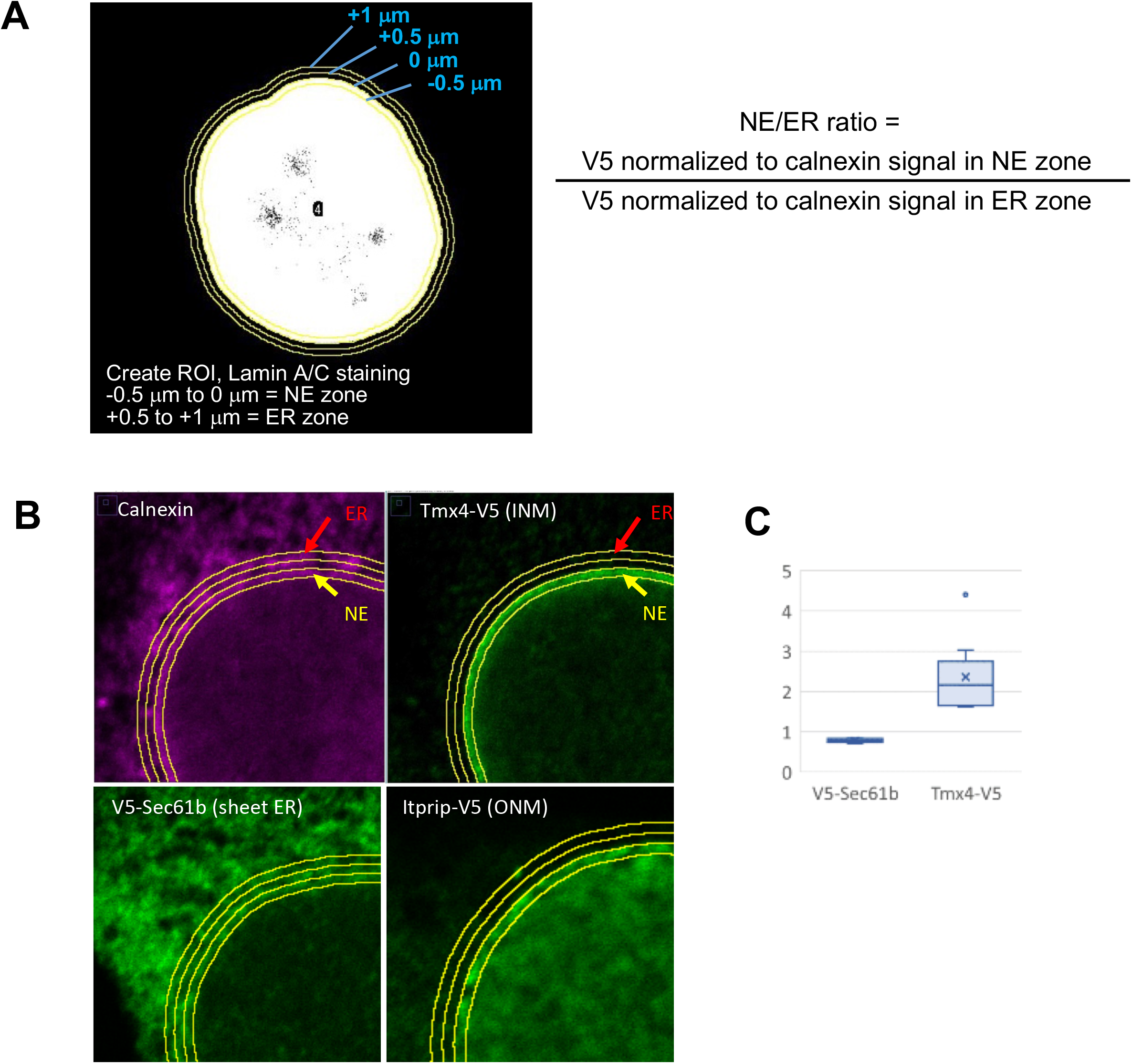
Scheme for quantification of the NE/ER concentration ratio of ectopically expressed proteins from immunofluorescent images. *A,* Left: Immunofluorescent labeling of cells to detect lamin A/C is used to create a region of interest (ROI), which is used to define 0.5 μm circumferential zones around the nuclear periphery containing the NE and peripheral ER for immunofluorescence quantification as depicted. Right: The NE/ER ratio of ectopic V5-tagged constructs is based on the V5 fluorescence intensity normalized to calnexin intensity in the corresponding zone. *B,* Top two panels show immunofluorescence labeling of endogenous calnexin and Tmx4-V5 (an INM marker) in the same cell, using the demarcation method in *(A)*. The bottom two panels show localization of ectopic Sec61*β* (a sheet ER marker) and Itprip (an ONM marker) with respect to zones defined as in *(A)*. C, Box and whisker plot showing quantification of the NE/ER concentration ratio for ectopic Sec61*β* and Tmx4 as in *(A)*.

**Fig. S4.**
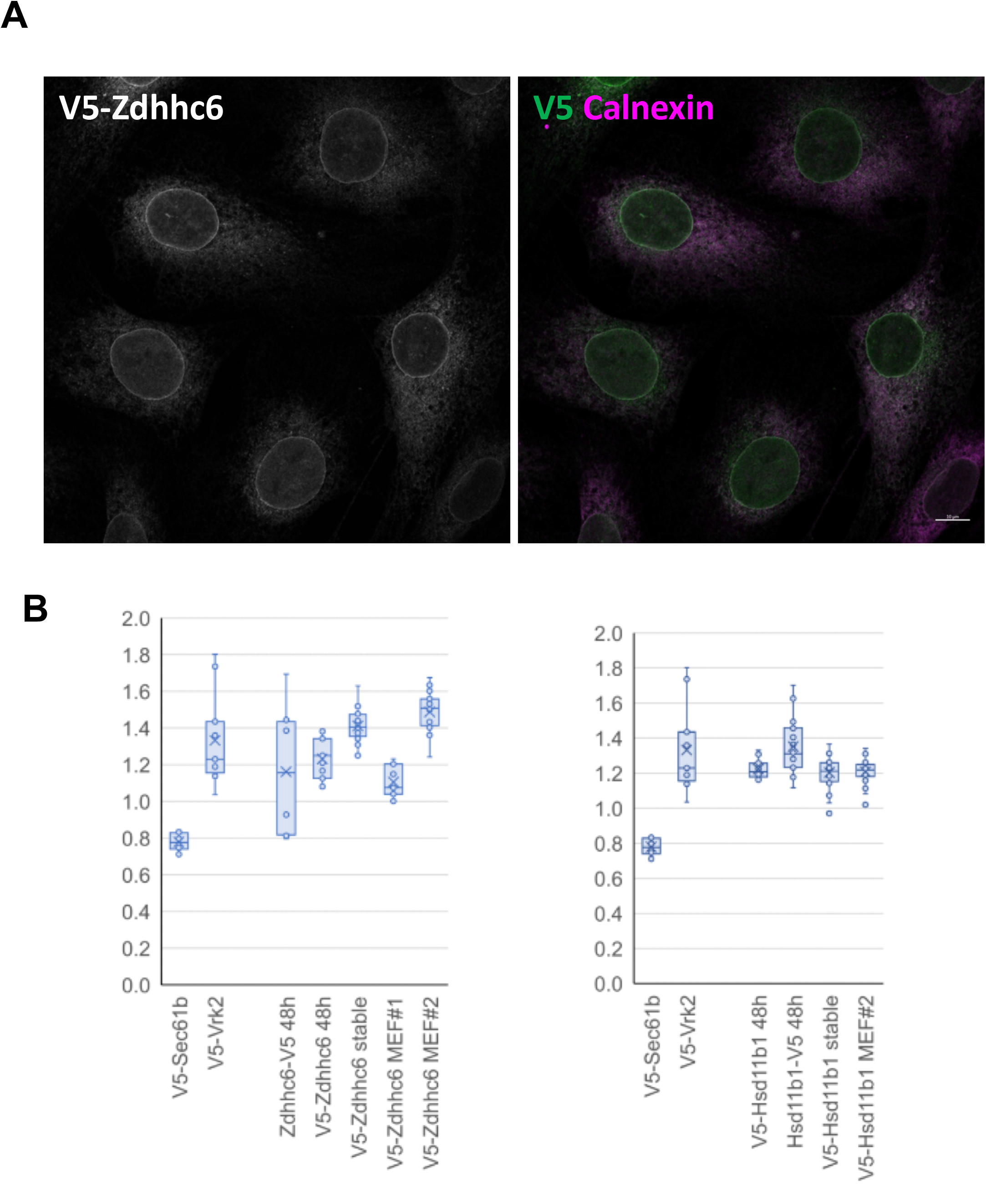
Comparison of NE/ER concentration ratio of Zdhhc6 and Hsd11b1 in different expression conditions and cell types. *A,* Micrographs showing a field of C3H cells labeled for immunofluorescence microscopy to detect transiently expressed Zdhhc6-V5 (at 48 hours after lentivirus infection) vs endogenous calnexin. *B,* Box and whisker plot depicting quantification of NE/ER ratio of Zdhhc6 (left graph) or Hsd11b1 (right graph) in transiently transduced cells, or in different stably transduced cell populations. The first four entries in each graph were taken from Fig. 3, bottom panel. The remaining entries are values calculated from different stably transduced C3H or MEF populations, as indicated. Note that V5-Zdhhc6 MEF population #1 was created with a several-fold higher titer of virus that MEF population #2, and likely contains cells with multiple lentiviral integrations.

**Fig. S5.**
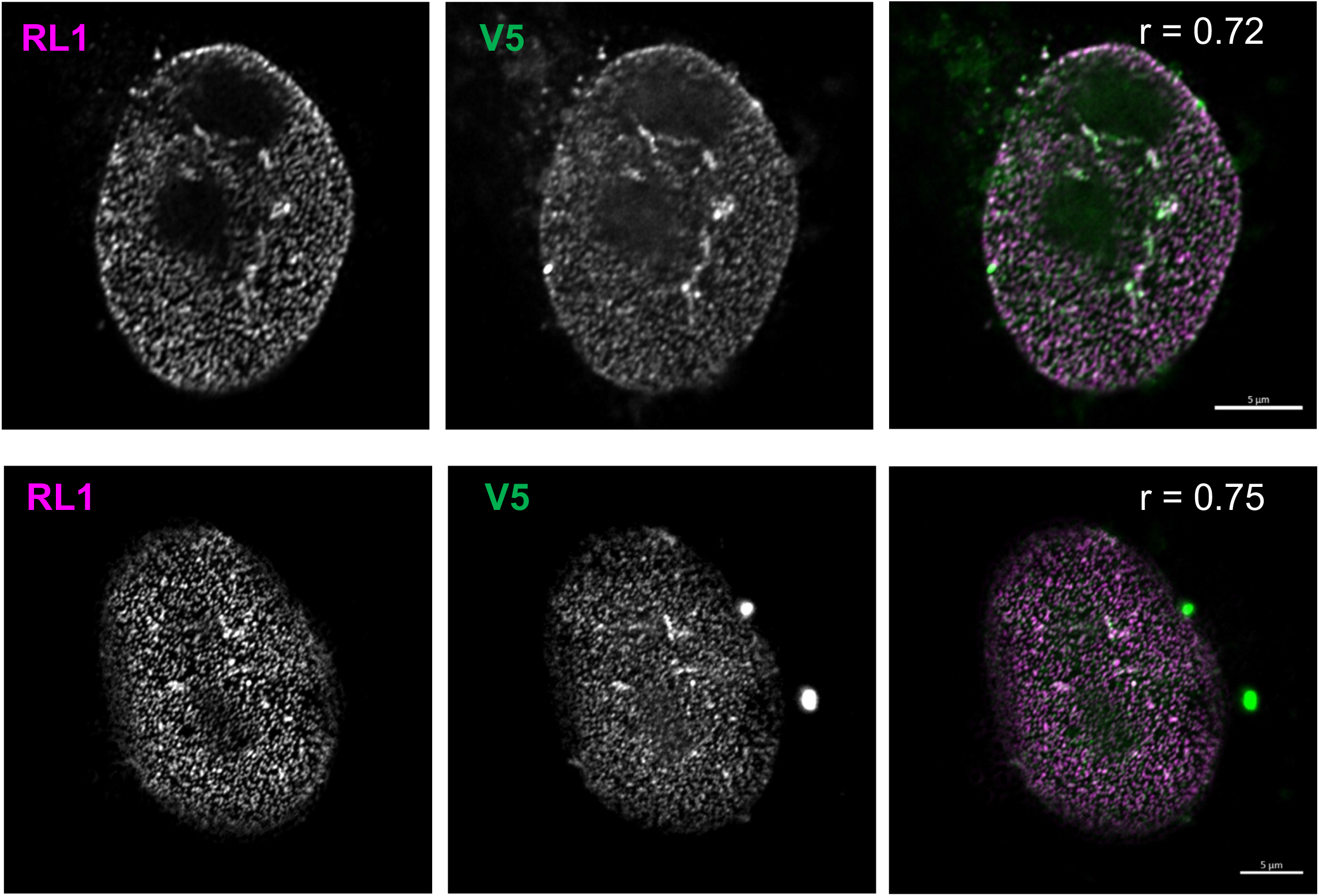
Comparison of the localization of ectopic Tmem209 and NPCs. Immunofluorescence micrographs of two MEFs analyzed 48 h after transient transduction with V5-Tmem209. Shown are Airyscan confocal images of the co-staining detecting the RL1 monoclonal antibody that recognizes FG repeat nucleoporins (RL1) and ectopic Tmem209 (V5). The merged images are in the rightmost panels. Pearson’s correlation coefficient comparing the RL1 and V5 images (r) is indicated for each cell, and reflect strong correlation. Bar = 5 μm.

**Fig. S6.**
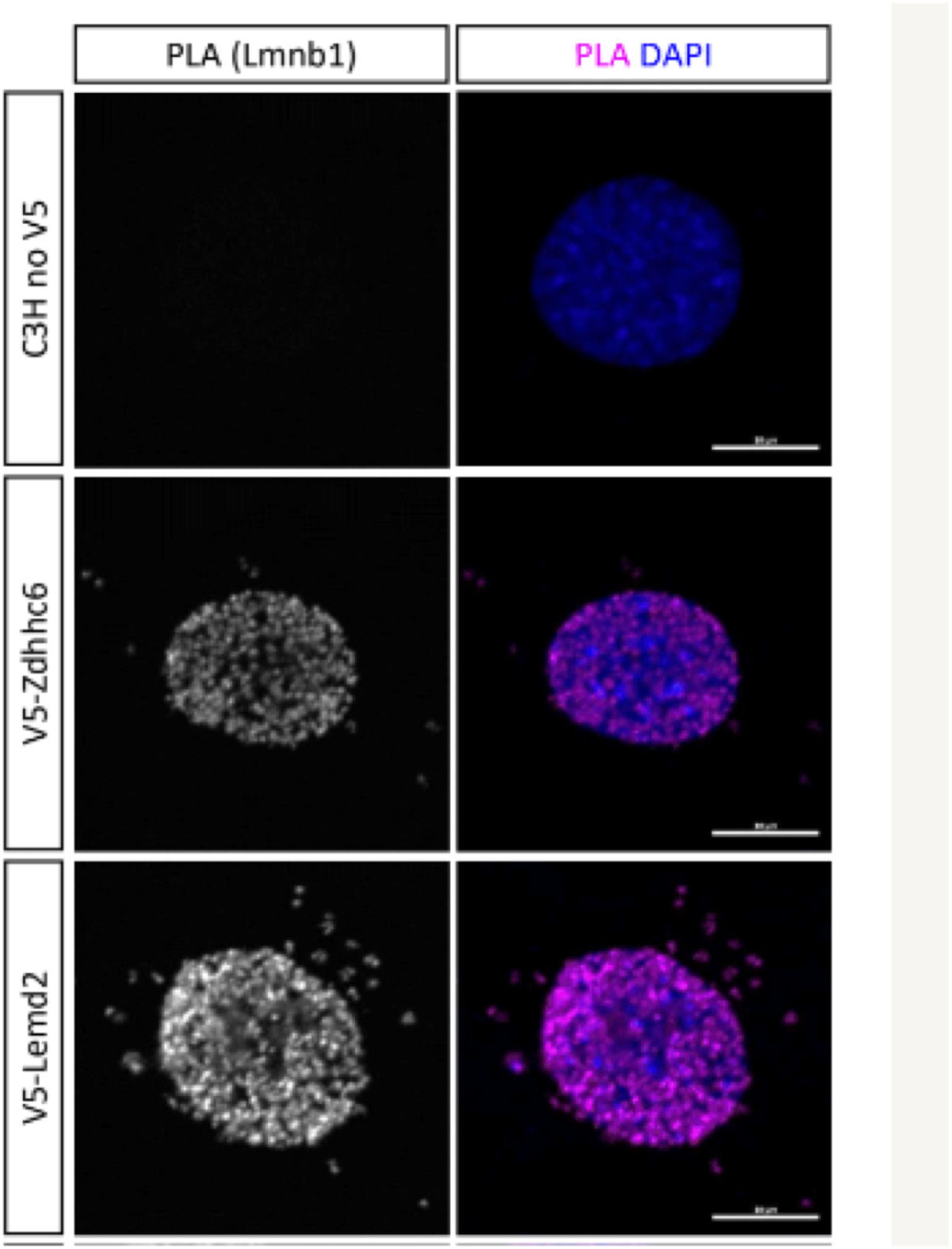
Localization of a pool of Zdhhc6 to the INM using the proximity ligation assay (PLA). Detection involved C3H cells stably transduced with Myc-tagged lamin B1 and transiently expressed Zdhhc6-V5 or Lemd2-V5 (a control INM protein). PLA signal, left panels; merge of PLA signal with DNA staining, right panels. Bar, 10 μm.

**Fig. S7.**
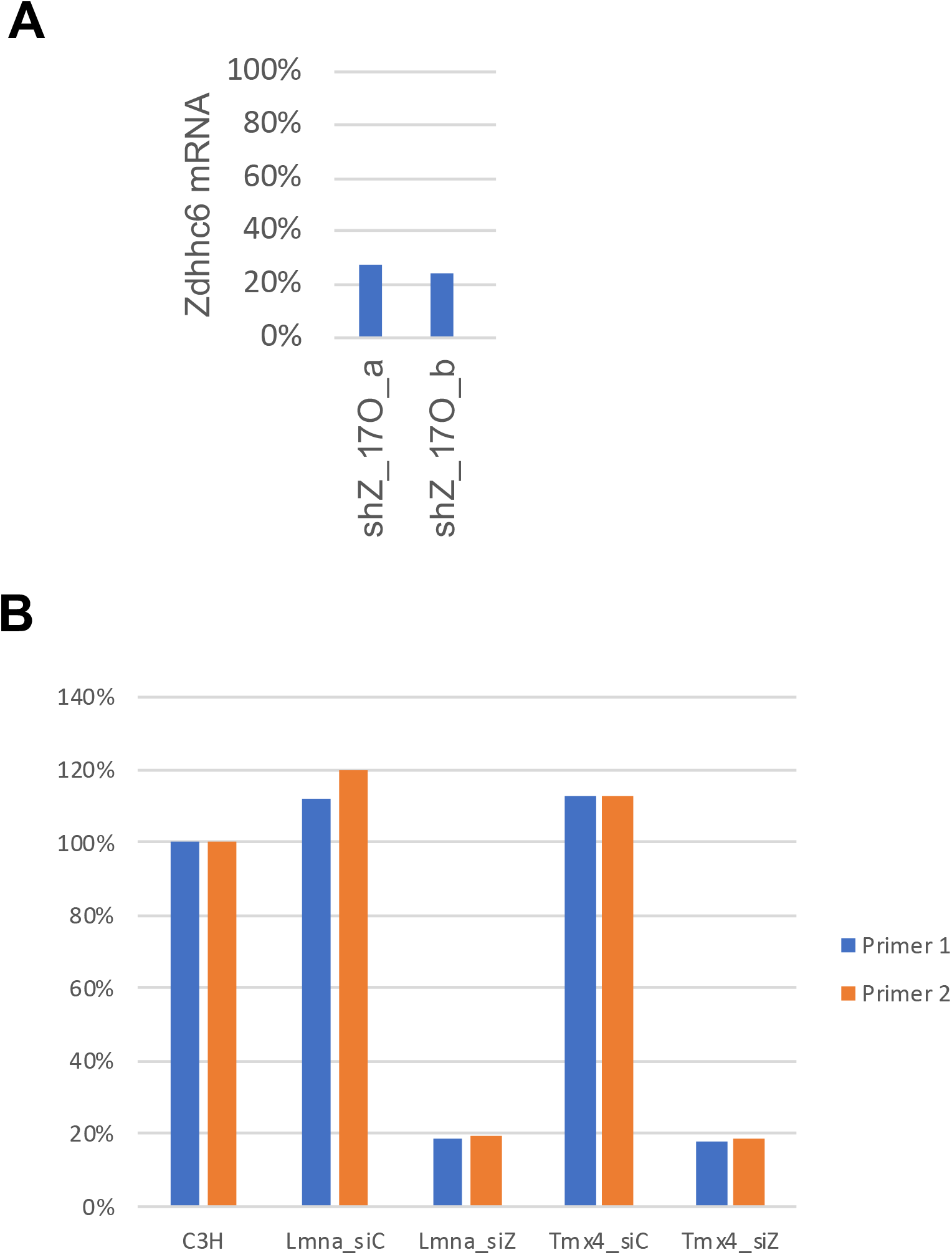
Depletion efficiency of Zdhhc6 mRNA in cells transduced with shRNA or transfected with siRNA. *A,* Levels of Zdhhc6 mRNA, as determined by qRT-PCR, in C3H cells stably transduced with an shRNA vector targeting Zdhhc6. Shown is analysis of cell samples “a” and “b” used for metabolic labeling with 17ODYA and quantification of palmitoylation changes with Zdhhc6 depletion (Supplemental Table S4 and Fig. 4B). *B,* Levels of Zdhhc6 mRNA, as determined by qRT-PCR, in various C3H cell populations transiently transfected with an siRNA pool targeting Zdhhc6 (siZ) or with a control siRNA pool (siC). The cell populations analyzed, as indicated, were WT C3H cells, or C3H cells that were stably transduced with vectors expressing V5-lamin A or Tmx4-V5. Proteins analyzed by immunofluorescence targeting assay.

